# Depletion of rRNA Methyltranferase METTL5 Enhances Anti-Tumor Immune Response via Neoantigen Generated from Cryptic Translation

**DOI:** 10.1101/2025.06.06.658288

**Authors:** Yangyi Zhang, Xiaoyan Shi, Yuci Wang, Ruiqi Wang, Folan Lin, Yanlan Cao, Wanqiu Li, Hao Chen

## Abstract

Tumor neoantigens play a pivotal role in eliciting tumor-specific immune responses and holds the promise for personalized immunotherapy. However, previous studies mainly focused on the tumor-specific neoantigens derived from genomic mutation and aberrant RNA splicing, limiting the repertoire of targetable neoantigens. Here, we demonstrate that inhibition of rRNA methyltransferase METTL5 translationally increases neoantigen production and enhances anti-tumor immunity. Mechanistically, METTL5-mediated m^6^A modification at the decoding center of small ribosomal subunit maintains the proper function of ribosome during mRNA translation. *METTL5*-deficiency decreases translation fidelity and increases production of tumor cell-specific antigens derived from non-canonical translation. Furthermore, we found that *Mettl5*-depletion increased CD8⁺T cell infiltration density and T cell receptor (TCR) repertoire diversity in murine tumor models. Importantly, this immunostimulatory effect strictly depended on intact antigen presentation pathways, suggesting that *Mettl5* knockout exerts its effects primarily through neoantigen generation. Together, this study uncovers the intrinsic mechanisms sustaining mRNA translation accuracy, elucidates a novel source of tumor neoantigen generation, and proposes a new strategy to enhance immunotherapy through targeting mRNA translation.

## INTRODUCTION

The complexity of cancer therapy originates from the high heterogeneity of tumors, dynamic immune escape mechanisms, and the multi-faceted challenges posed by the immunosuppressive tumor microenvironment^1–8^. Conventional therapies (e.g., chemotherapy and radiotherapy) often cause severe toxic side effects with limited therapeutic efficacy, whereas immunotherapy, despite its advantages of low toxicity and durable responses, remains constrained by low response rates and the development of acquired resistance^3–5,9–16^.

A promising solution lies in the identification of tumor-specific mutations to generate neoantigens, which can activate polyclonal T-cell responses, overcome immune tolerance, and avoid off-target toxicity^17–24^. This “dynamic targeting” approach offers a means to address heterogeneity and drug resistance^4,5,25,26^. Neoantigens, derived from short peptides (8–25 amino acids) resulting from errors in gene expression regulation, are presented on tumor cell surfaces by human leukocyte antigen (HLA) molecules, enabling T-cell recognition^27–30^. Unlike self-antigens, neoantigens are not subject to central tolerance mechanisms and exhibit robust immunogenicity, making them ideal targets for immunotherapy^17,18,21,23,31–33^.

Certain tumor mutations had been found to enhance the quality and quantity of functional neoantigens, such as those targeting highly immunogenic mutations like KRAS G12D, IDH1 R132H, and TP53^34–37^. These neoantigen vaccines have demonstrated the ability to activate T-cell responses in solid tumors such as pancreatic cancer and even penetrate the blood-brain barrier in glioma^38,39^. Furthermore, the combination of personalized vaccines (e.g., NeoVax) with immune checkpoint inhibitors has advanced clinical exploration in precision immunotherapy, suggesting that the quality of neoantigens generated by tumor mutations may guide therapeutic efficacy^40^.

Despite recent advances in tumor neoantigen, clinical application of neoantigen-targeted therapies faces multiple challenges. First, most antigens arising from tumor mutations are of low quality (e.g., synonymous mutations or cryptic epitopes), exhibiting insufficient immunogenicity^41–45^. Second, certain mutated peptides share high sequence homology with self-proteins, leading to weakened anti-tumor immune responses due to central tolerance (thymic selection) or peripheral tolerance (T cell exhaustion) mechanisms^16,46^. More critically, the process of immune editing within the tumor microenvironment continuously eliminates highly immunogenic epitopes, leaving residual antigen pools dominated by low-affinity variants that further impair T cell activation efficiency^43,47^. To overcome these limitations, current intervention strategies-such as CRISPR/Cas9-induced mutations or DNA-damaging agents to enhance neoantigen load, which can transiently expand neoantigen numbers^5,48,49^. However, their irreversible genomic modifications risk exacerbating tumor heterogeneity or causing normal tissue toxicity, significantly hindering clinical application of these approaches^50,51^.

To address the critical challenge of limited availability of tumor-specific neoantigens, we explored translational regulation as a reversible and safer strategy to augment neoantigen production. In current study, we integrated multi-omics analysis with functional validation to demonstrate that loss of METTL5-mediated m^6^A modification in ribosomes induces neoantigen generation through disrupting translational fidelity. The neoantigens derived from mistranslation significantly diversified the T cell receptor (TCR) repertoire and enhanced CD8⁺ T cell infiltration. Our findings expand both the diversity and sources of tumor neoantigens, offering a novel pathway to enhance anti-tumor immune responses.

## RESULTS

### METTL5 is identified as a negative regulator of tumor immunogenicity

To identify the chemical modifications of ribosomal RNA that relate with resistance to immunotherapy, we systematically compared the differential expression profiles of 11 key rRNA modifiers between responders (n=82) and non-responders (n=72) of ICI (Immune-checkpoint inhibitors)-treated samples^52^(Fig. 1A). Notably, the result revealed that the expression level of METTL5 is negatively associated with immunotherapy responses (Fig. 1A). Previous studies demonstrated that METTL5 must form a heterodimeric complex with TRMT112 to functionally catalyze m^6^A methylation of adenosine at position 1832 in 18S rRNA^53–55^. Interestingly, elevated TRMT112 expression also serves as a negative predictor of immunotherapy response, which suggests that 18S rRNA m^6^A modification mediated by METTL5-TRMT112 may be intrinsically linked to tumor immunity.

**Fig. 1:**
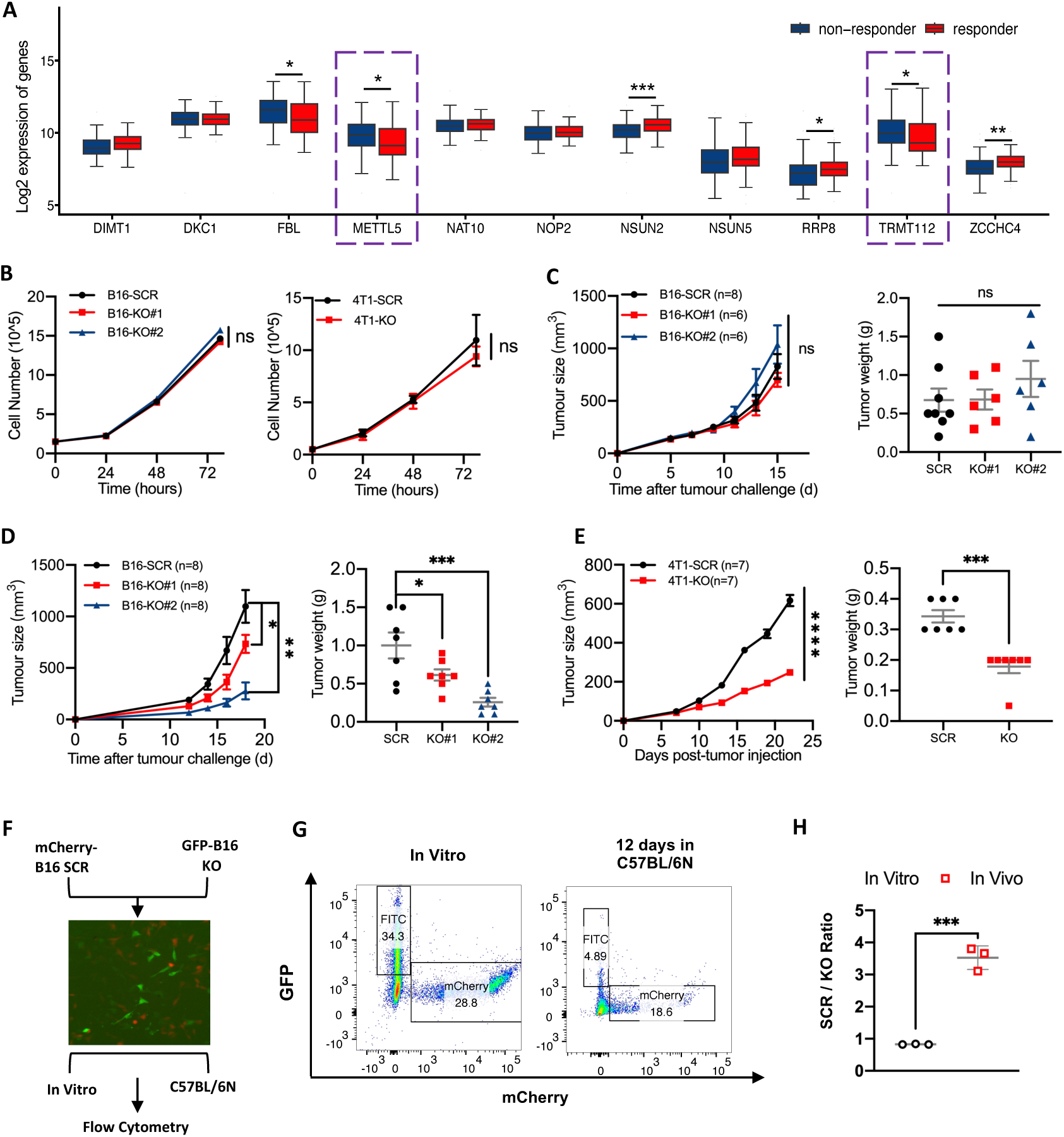
*METTL5* depletion attenuates tumour growth. **A**, Relationship between the expression levels of ribosomal RNA modifiers and response to immune checkpoint blockade (ICB) therapy. METTL5 and TRMT112 are highlighted with purple dashed boxes. **B**, *In vitro* cell growth assay of SCR and *Mettl5*-KO cells in B16-F10 (left) and 4T1 (right) cell lines. **C**, Growth curve of tumors in nude mice subcutaneously implanted with B16-F10 SCR or *Mettl5* deficiency cells. The tumor growth (left) and final tumor weights (right) were compared. **D**, Growth curve of tumors in C57BL/6N immunocompetent mice subcutaneously implanted with B16-F10 SCR or *Mettl5*-KO cells. The tumor growth (left) and final tumor weights (right) were compared. **E**, Growth curve of tumors in Balb/c immunocompetent mice subcutaneously implanted with 4T1 SCR or *Mettl5*-KO cells. The tumor growth (left) and final tumor weights (right) were compared. **F-H**, *In vivo* competition assay between mCherry-labeled SCR control and GFP-labeled *Mettl5*-KO B16-F10 cells. Schematic diagram of the competitive growth assay (**F**). mCherry-labeled SCR and GFP-labeled-KO cells were mixed at a 1:1 ratio and either subcutaneously injected into C57BL/6N mice or co-cultured *in vitro*. Cell ratios were quantified by flow cytometry on Day 12. Representative flow cytometry plots showing the composition of SCR and *Mettl5*-KO cells (**G**). Quantitative analysis of cell population ratios (**H**). **Note: A**, Data retrieved from NCBI Gene Expression Omnibus (GEO) were presented as median (IQR 25-75%, n=154). Student’s t-tests. **B**, two-way ANOVA. **C**, **D**, two-way ANOVA (left) and One-way ANOVA with Tukey’s test (right). **E**, two-way ANOVA (left) and Student’s t-tests (right). **H**, Student’s t-tests. **B**, **H**, Data were represented as mean ± SD (n=3). **C**, **D**, **E**, Data were presented as mean ± SEM. \**P*<0.05, \*\**P*<0.01, \*\*\**P*<0.001, ns: not significant.

To evaluate the regulatory role of METTL5 in tumor immunogenicity, we generated *Mettl5*-knockout B16-F10 melanoma and 4T1 breast cancer cell lines using CRISPR/Cas9-mediated gene editing (Fig. S1A). Sanger sequencing confirmed biallelic frameshift mutations at the *Mettl5* locus, with indel events clustered around the gRNA target site (Fig. S1A). Quantitative Reverse Transcription PCR (RT-qPCR) demonstrated significant transcript depletion (>70%) in both *Mettl5*-KO cell lines (Fig. S1B). Consistently, high-performance liquid chromatography-mass spectrometry (HPLC-MS) quantification revealed significant decrease of m^6^A modification in *Mettl5*-KO cells (Fig. S1C). Furthermore, cell growth assays showed no significant differences between *Mettl5*-KO and scrambled (SCR) control in either cell line (Fig. 1B). Similarly, clonogenic survival assays demonstrated comparable colony-forming abilities (Fig. S1D, S1E). These findings suggest that the loss of *Mettl5* does not significantly impair the proliferation capacity of tumor cells *in vitro*.

Then we established syngeneic tumor models to determine the immunological consequences of *Mettl5* depletion using immunocompromised (nude) and immunocompetent (C57BL/6N or Balb/c) mice, respectively. Consistent with the *in vitro* results, *Mettl5* loss did not influence the ability of B16-F10 cells to form tumors in nude mice with deficient immune systems (Fig. 1C). In contrast, when *Mettl5-*deficient cells were inoculated into syngeneic mouse hosts, their abilities to form tumors were significantly attenuated compared with control cells (Fig. 1D, E). To more precisely assess the role of METTL5 in tumor immunity, we carried out competitive cell growth assays to compare the proliferation rates of GFP-labeled *Mettl5*-KO and mCherry-labeled SCR B16-F10 cells under identical conditions. Consistently, the results demonstrated that *Mettl5*-deficient cells only exhibited growth suppression compared to control cells in immunocompetent mice (Fig. 1F-H). Collectively, these findings demonstrate that METTL5 regulates tumor growth through an immune-dependent mechanism.

### *Mettl5* knockout promotes immune cell infiltration in tumor

To elucidate how *Mettl5* depletion suppresses tumor growth specifically in immunocompetent hosts, we systematically characterized immune cell dynamics within the tumor microenvironment (TME). Immunohistochemical (IHC) analysis of B16-F10 and 4T1 tumors revealed significantly enhanced CD8^+^ T cell infiltration in *Mettl5*-KO tumor model compared to SCR control (Fig. 2A, B). To quantitatively profile immune populations, we isolated leukocytes from B16-F10 tumors using Percoll gradient centrifugation for multicolor flow cytometry analysis. *Mettl5*-KO tumors exhibited marked increases in CD8^+^ T cells, accompanied by elevated proportions of NK cells and TH cells (Fig. 2C, D). These findings were further supported by spleen immunophenotyping, which revealed expanded cytotoxic T lymphocyte (CTL) populations in *Mettl5*-KO tumor-bearing mice (Fig. 2E, F). Lastly, we performed single-cell RNA sequencing (scRNA-seq) to characterize infiltrating lymphocyte populations in freshly isolated tumor tissues (Fig. S2A). Unsupervised clustering identified major immune subsets including CD8^+^ T cells, B cells, macrophages, and dendritic cells (Fig. S2B-D). Notably, the TMEs of *Mettl5*-KO tumor contained higher proportions of CD8^+^ T cells compared to controls (4% vs 2.5%) (Fig. S2E). Therefore, scRNA-seq data consistently validated the immunostimulatory phenotype observed through IHC and flow cytometry, reinforcing METTL5’s regulatory role in tumor-immune interactions.

**Fig. 2:**
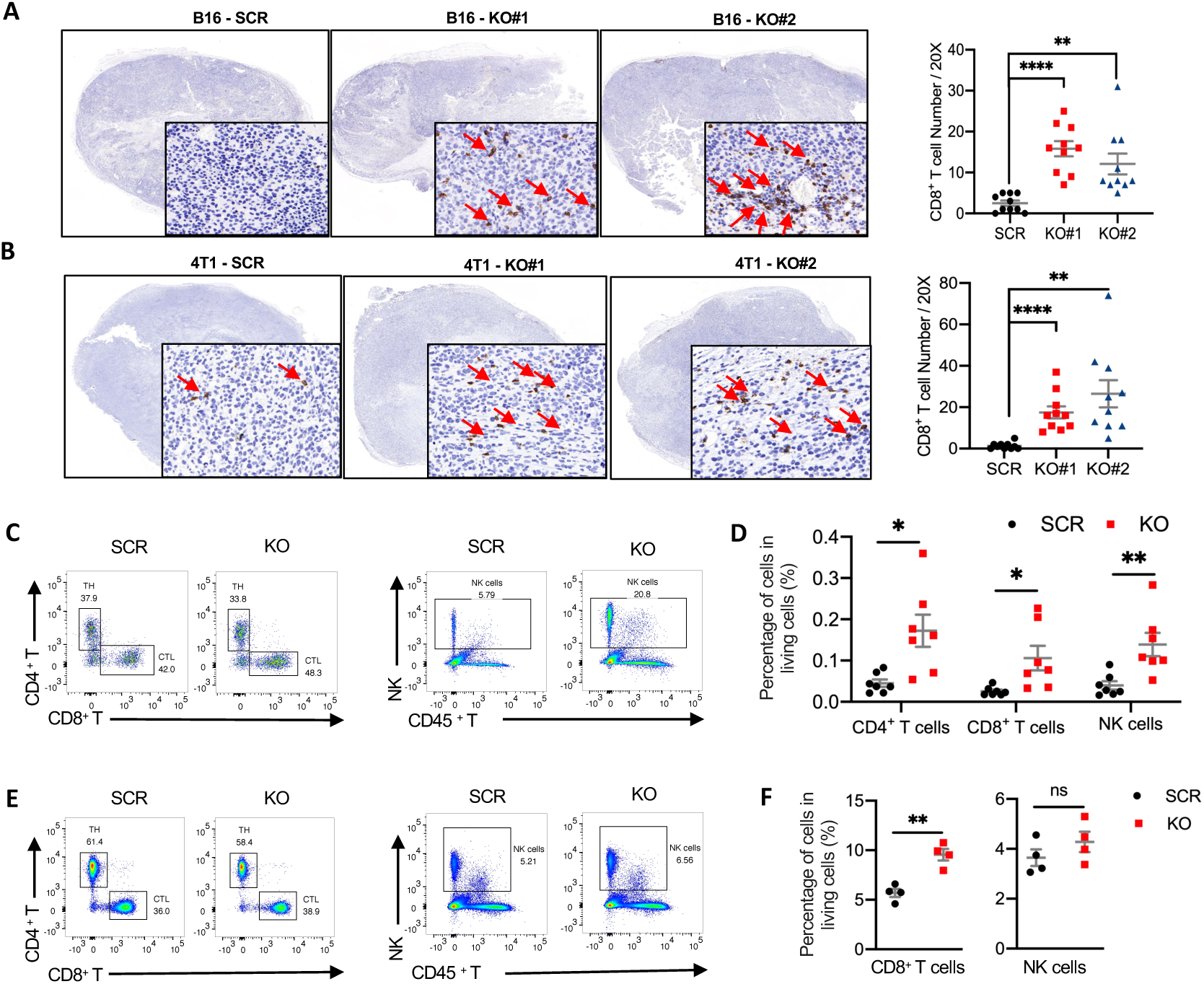
*Mettl5* deficiency enhances tumor immune cell infiltration. **A**, **B**, Immunohistochemistry (IHC) detected CD8^+^ cells in *Mettl5*-KO tumor tissues derived from B16-F10 (**A**) and 4T1 (**B**) models. Representative images (left) and quantified CD8^+^ cell counts (right) are shown. 10 random 20× fields per sample were analyzed. CD8^+^ cells are indicated by red arrows. **C-F**, Flow cytometry analysis of immune cell populations in the tumor microenvironment (**C**) and spleen (**E**) of C57BL/6N mice inoculated with SCR or *Mettl5*-KO B16-F10 cells, and representative flow cytometry image showing the distribution of CD4^+^ T cells, CD8^+^ T cells (left) and NK cells(right). The relative frequency of immune cells amongst living cells from tumors (**D**) and spleen (**F**) were assessed. **Note: A**, **B**, One-way ANOVA with Tukey’s test. **D**, **F**, Student’s t-tests. **A**, **B**, **D**, **F**, Data were represented as mean ± SD. \**P*<0.05, \*\**P*<0.01, \*\*\*\**P*<0.0001, ns: not significant.

### Loss of *Mettl5* enhances the response to immune checkpoint blockade

Given that *Mettl5* knockout has been shown to enhance tumor-infiltrating capacity of CD8^+^ T cells, which play a critical role in antitumor immunity^56^, we further asked whether depleting *Mettl5* synergizes with immune checkpoint blockade. In C57BL/6N mice bearing B16-F10 melanomas, *Mettl5*-KO tumors exhibited significantly reduced growth compared to SCR controls following anti-PD-1 treatment (Fig. 3A). Importantly, the combination of *Mettl5* depletion and anti-PD-1 therapy demonstrated the most robust anti-tumor effect, with median survival prolonged by 3 days compared to groups only administrated to anti-PD-1 antibody (Fig. 3A). Furthermore, this finding was also recapitulated in the PD-1-resistant 4T1 breast cancer model. *Mettl5* loss significantly boosted antitumor immune responses, including reduced tumor volumes, and extended survival compared to other groups (Fig. 3B).

**Fig. 3:**
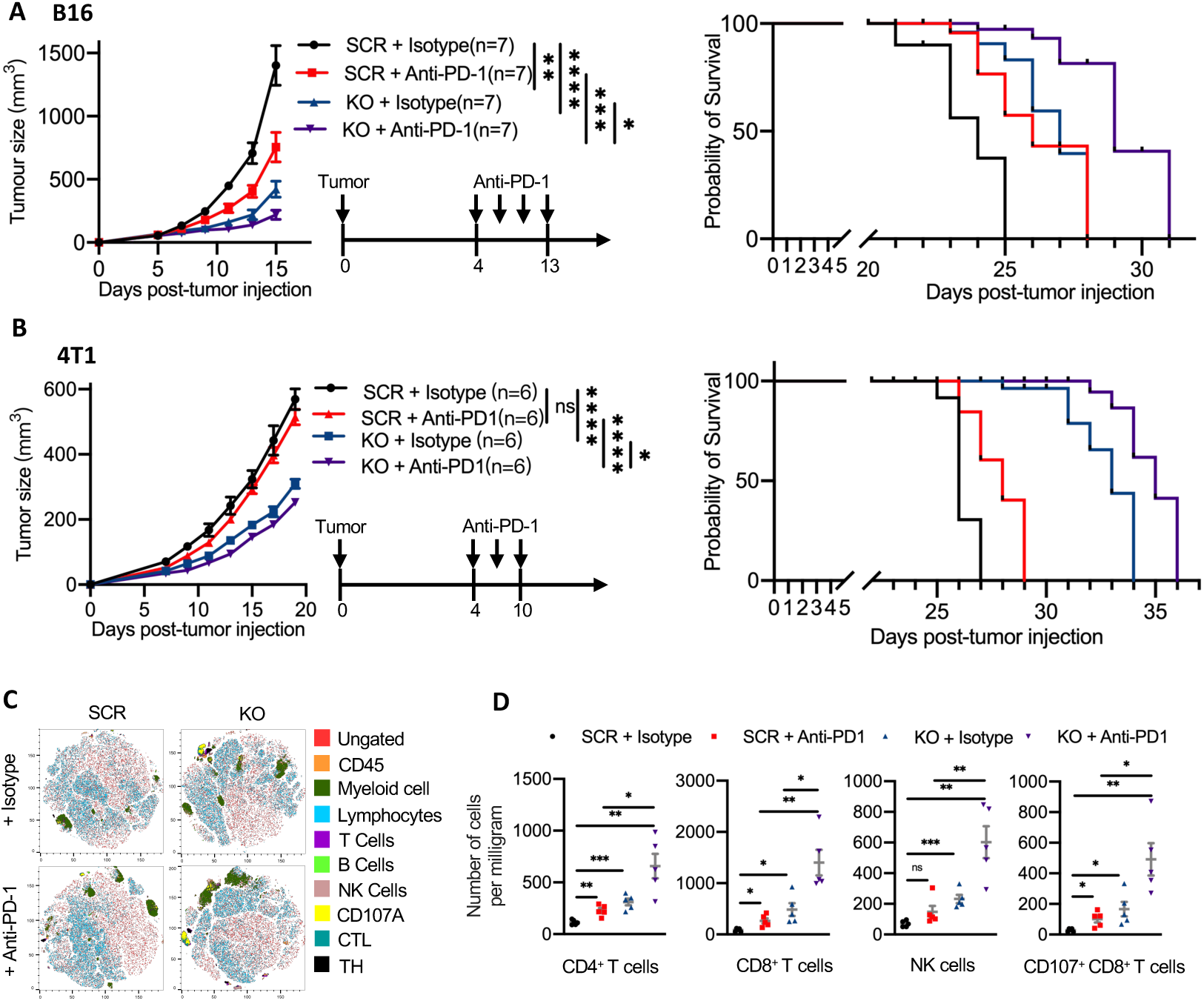
Depletion of *Mettl5* increases the efficacy of tumor immunotherapy. **A**, Anti-PD-1 antibody was administered to *Mettl5*-KO and SCR B16-F10 tumor-bearing C57BL/6N mice on days 4, 7, 10, and 13 post-implantation, followed by longitudinal monitoring of tumor volume (left) and overall survival (right). **B**, Anti-PD-1 antibody was administered to *Mettl5*-KO and SCR B16-F10 tumor-bearing Balb/c mice on days 4, 7, and 10 post-implantation, followed by longitudinal monitoring of tumor volume (left) and overall survival (right). **C**, t-SNE analysis of immune cell populations in the tumor microenvironment of tumor in C57BL/6N mice. Mice were treated with either isotype control or anti-PD-1 antibody. The analysis includes the following immune cell populations: Lymphocytes, CD45⁺ cells, Myeloid cells, T Cells, B Cells, NK Cells, CD107a⁺ cells, CTL, and T Helper (TH) cells. The plots show the distribution of these immune cell populations in the tumor microenvironment for SCR and KO groups treated with isotype control or anti-PD-1 antibody. **D**, Quantitative analysis of immune cell populations in the tumor microenvironment of tumor in C57BL/6N mice. The number of CD4^+^ T cells, CD8^+^ T cells, NK cells, and CD107a^+^ CD8^+^ T cells per milligram of tumor tissue was determined by flow cytometry. **Note:** Mouse survival curves were plotted using endpoints: death or tumor volume exceeding 2000 mm³. **A**, **B**, two-way ANOVA. Data were presented as mean ± SEM. **D**, Student’s t-tests. Data were represented as mean ± SD (n=5). \**P*<0.05, \*\**P*<0.01, \*\*\**P*<0.001, \*\*\*\**P*<0.0001, ns: not significant.

To further evaluate the synergic effect of *Mettl5* depletion and anti-PD-1 blockade, we analyzed tumor-infiltrating immune cells in B16-F10 melanoma tissues using flow cytometry. Compared to SCR tumors, *Mettl5*-KO tumors exhibited significantly higher proportions of CD8^+^ T, CD4^+^ T cells, and NK cells among intratumor cells (Fig. 3C-D, Fig. S3A-B). Previous studies indicated that anti-PD-1 therapy expanded tumor-infiltrating cytotoxic immune cell populations, including CD8^+^ T cells, and augmented their functional activity (as measured by elevated CD107a expression)^57^. Notably, *Mettl5*-KO synergized with anti-PD-1 treatment indeed enhanced these effects further (Fig. 3D, S3B, rightmost panel). Together, our results revealed that *Mettl5* knockout potently augments the anti-tumor efficacy of anti-PD-1 treatment, suggesting that METTL5 inhibition can overcome immune checkpoint inhibitor resistance and improve therapeutic outcomes in cancer immunotherapy.

### *Mettl5* depletion decreases ribosomal translation fidelity

Recent advances have highlighted the critical role of RNA modifications in ribosome function and translation regulation^58,59^. Though METTL5 had been identified as the primary methyltransferase responsible for catalyzing m⁶A1832 modification on 18S rRNA^60–62^, its precise molecular functions, particularly regarding its roles in translation regulation under physiological and pathological conditions, remain poorly understood. To uncover its functional roles, we first investigate the individual 3D environments of m⁶A1832 modification inside the ribosome. As shown in Fig. 4A-B, the m^6^A1832 site is located within the core region of the 40S subunit’s decoding center, specifically at the h44-h45 helical junction of 18S rRNA. Adjacent nucleotides, including A1831, C1704, Cm1703, and G1702, stabilize the conformation of the modification site through base stacking and hydrogen bonding (Fig. 4A). Additionally, these residues directly interact with ribosomal proteins uS12, uS11, and uS5, which are known to play pivotal roles in regulating the dynamics of the decoding center ^63,64^. Specifically, uS12 and uS11 are involved in the precise recognition of mRNA codons by interacting with the anticodon loop of tRNA^65–67^. These findings suggest that m⁶A1832 methylation may enhance the rigidity of h44 (a central helix in the decoding center) or optimize local nucleotide-protein interactions, thereby reducing conformational fluctuations during translation and improving ribosome proofreading capabilities to ensure translational fidelity. Interestingly, the methylation enviroment of A1832-C1702 pair is conserved from bacteria to mammal. In *E. coli*, although A1500 (corresponding A1832 in human) is unmodified, its paired C1402 (C1702 in human) carries an m⁴Cm modification (Fig. 4C), which is critical for maintaining the small ribosomal subunit rRNA structure, and enhancing translation fidelity^68,69^.

**Fig. 4:**
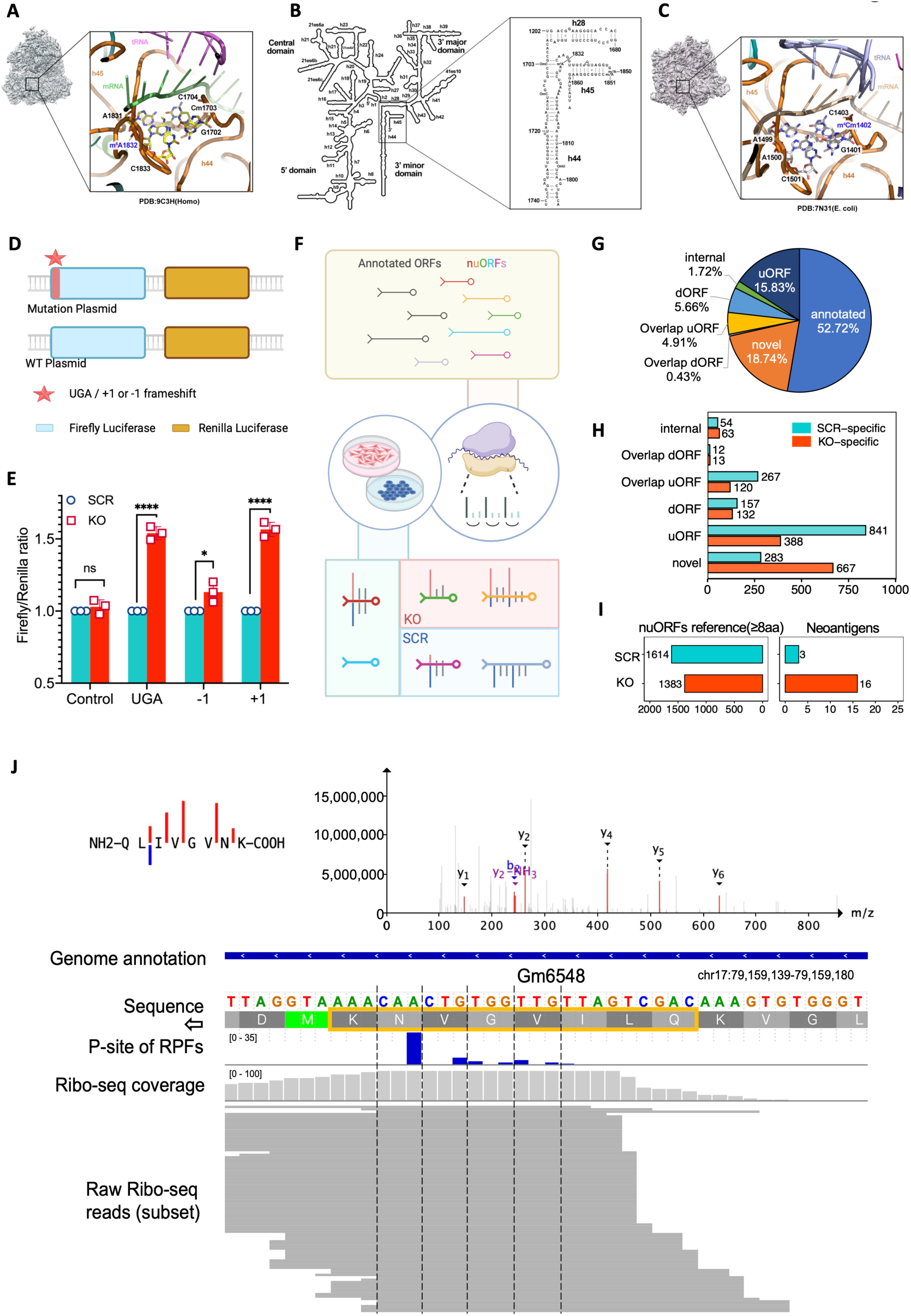
Depletion of *Mettl5* induces neoantigen generation. **A**, Close-up view of the human 40S ribosomal subunit decoding center highlighting the m⁶A1832 modification (blue). **B**, Overall structure of 18S rRNA within the decoding center. **C**, Close-up of the *E. coli* 30S ribosomal subunit decoding center featuring the m⁴Cm1402 modification (blue). **D**, Scheme of reporter vector. Dual-luciferase plasmid constructs were designed with a UGA stop codon, +1 frameshift, or −1 frameshift upstream of the firefly luciferase gene, with Renilla luciferase as an internal control. **E**, Translation efficiency of wildtype and mutant firefly luciferases in SCR and *Mettl5* KO cells was revealed by a reporter assay. Firefly luciferase activity was normalized to Renilla luciferase, with the SCR group set as the baseline (relative ratio = 1). **F**, Schematic workflow of generating nuORFdb v.1.0. ORFs from ribosome profiling (Ribo-seq) data were classified into annotated ORFs and novel ORFs (nuORFs), which were compiled as a database for subsequent analyses. For differential nuORF expression between *Mettl5*-KO and SCR groups, we classified nuORFs into three categories: (1) Shared nuORFs (expressed in both groups with <1.5-fold TPM difference), (2) *Mettl5*-KO-specific nuORFs (exclusive to *Mettl5*-KO or with ≥1.5-fold higher TPM than SCR), and (3) SCR-specific nuORFs (exclusive to SCR or with ≥1.5-fold higher TPM than *Mettl5*-KO). **G**, Pie chart illustrating the classification of nuORFs identified in *Mettl5*-KO and SCR group. The chart shows the percentage distribution of different nuORF categories relative to the total nuORFs. **H**, Differential analysis of nuORF categories between *Mettl5*-KO and SCR groups. **I**, Mass spectrometry-based identification of MHC-bound peptides in SCR and *Mettl5*-KO samples, nuORF-derived neoantigens were identified using corresponding nuORFs (>8 aa) as reference sequences. **J**, A representative neoantigen (NH2-Q-L-I-V-G-V-N-K-COOH) identified by MHC-I immunopeptidomics and Ribo-seq in *Mettl5*-KO tumors. The characteristic 3-nucleotide periodicity of ribosome-protected fragments confirmed its translational activity. Top panel: the mass spectra for the identified neoantigen; Bottom panel: Ribo-seq revealing nuORF reading frames. Blue bars, in-frame reads; LC-MS/MS-detected peptide was highlighted with a yellow box. **Note: E**, Student’s t-tests. Data were represented as mean ± SD (n=3). \**P*<0.05, \*\*\*\**P*<0.0001, ns: not significant.

Based on the above structural and evolutionary evidences, we speculate that METTL5 is likely involved in controlling mRNA translation fidelity. To test this hypothesis, we set up a dual-luciferase reporter system to assess the accuracy of mRNA translation upon loss of *Mettl5* (Fig. 4D). In the control plasmids (without pre-stop/frameshift mutations), no significant difference was observed in the Firefly/Renilla ratio between SCR and KO cells, indicating that *Mettl5* knockout does not affect the basal translation efficiency (Fig. 4E). However, the synthesis of mutant Firefly luciferase in *Mettl5*-KO cells was significantly higher than that in SCR cells (Fig. 4E). Specifically, the UGA stop codon group demonstrated increased ribosome readthrough of termination signals, and the −1/+1 frameshift mutation groups revealed a reduced ability of ribosomes in maintaining protein synthesis reading frame (Fig. 4E). These results directly support the notion that *Mettl5* knockout decreases ribosomal translation fidelity, manifesting as enhanced tolerance to termination signal readthrough and frameshift errors, thereby promoting the expression of aberrant proteins encoded by mutant plasmids.

### *Mettl5* depletion induces immunogenic neoantigen production by promoting cryptic translation

Given the observed translational perturbations in *Mettl5*-KO cells, we hypothesized that the impaired translational fidelity potentially induce the cryptic translation of Novel or Unannotated Open Reading Frames (nuORFs). This could promote tumor neoantigen generation and consequently enhance tumor immunogenicity^70,71^. To test this hypothesis, we performed hierarchical prediction of translated nuORFs in *Mettl5*-KO versus SCR tumor cells using ribosome profiling (Ribo-seq) (Fig. 4F, S4A, S4B). The actively translated ORFs were categorized into annotated ORFs and nuORFs, and nuORFs were further classified into shared nuORFs, KO-specific nuORFs (nuORFs with TPM values 1.5-fold higher in the KO than SCR), and SCR-specific nuORFs. Subsequently, all nuORFs were categorized into five subtypes based on previous study^71^: novel ORFs, uORFs (upstream ORFs), dORFs (downstream ORFs), overlap_uORFs (overlapping upstream ORFs), and overlap_dORFs (overlapping downstream ORFs) (Fig. 4G). Further analysis revealed that nuORFs primarily originate from untranslated regions of protein-coding genes, pseudogenes, and long non-coding RNAs (lncRNAs) (Fig. S4C). In contrast to annotated ORFs, considerable portion of nuORF translation initiation is heavily dependent on non-canonical start codons (particularly CUG) (Fig. S4D), implying active translation in putative non-coding genomic regions. Additionally, the polypeptides translated from nuORFs are also significantly shorter than those produced by annotated ORFs (Fig. S4E). Importantly, we found that though *Mettl5* knockout had minimal impact on translation efficiency of annotated ORFs (Fig. S4F). Interestingly, *Mettl5* knockout significantly enhanced the translation of novel nuORFs among the five subtypes of nuORFs (Fig. 4H). Moreover, further analysis of reading frame integrity revealed that in the annotated ORF group, the SCR group had a higher in-frame rate than the KO group (Fig. S4G, left), suggesting an increased probability of translation frameshifts of the annotated ORFs in the KO group. As to the nuORF, the KO group exhibited a significantly higher in-frame rate than the SCR group (Fig. S4G, right), indicating that the extent of translation activities in non-coding regions are much higher in the KO than SCR group.

To verify whether the translated nuORFs have the potential to become neoantigens presented by MHC-I, we confirmed the existence of nuORF-derived neoantigens through MHC-I immunopeptidomics. Based on the property that neoantigens require a polypeptide length of at least 8 amino acids, we selected nuORFs with amino acid lengths no less than 8 as the reference database for the tandem mass spectral (MS/MS) library search. Although the number of KO-specific nuORFs was less than that in the SCR group, MS/MS search detected 16 neoantigens derived from nuORFs in KO group, significantly higher than the number (only 3) of nuORF-derived neoantigens detected in the SCR group (Fig. 4I, J). Together, these findings indicate that *Mettl5* loss enhances production of neoantigens generated by mistranslation of nuORFs.

### *Mettl5* deficiency increases TCR clonal diversity in tumors

Considering that *Mettl5*-deficient tumors increased neoantigen loads, we hypothesized that *Mettl5* knockout would drive enhanced T-cell receptor (TCR) repertoire diversity in tumor-bearing hosts. Therefore, we performed TCR sequencing on peripheral blood samples from B16-F10 melanoma mouse models at experimental endpoints and conducted comparative TCR repertoire analysis between *Mettl5*-KO and SCR groups. The CDR3 length distribution histograms revealed conserved patterns between groups, with dominant peaks observed at 10–15 amino acid (aa) lengths (Fig. S5A). Quantitative analysis showed obvious differences in TCR repertoire diversity between *Mettl5*-KO and SCR tumor-bearing mice (Fig. 5A). The Shannon entropy of the KO group (mean = 11.5) was significantly higher than that of the SCR group (mean = 10.5, p < 0.05), indicating greater TCR repertoire diversity in tumor-bearing hosts with *Mettl5*-deficient tumors (Fig. 5A, left). Furthermore, the clonality index was significantly lower in the KO group compared to the SCR group, suggesting reduced dominance of private clones (Fig. 5A, right) and a more evenly distributed clonal landscape in *Mettl5*-deficient tumor-bearing mice (Fig. 5B). Consistently, the diversity estimations across Hill diversity profiles (Q values 0–6) demonstrated higher diversity indices in the KO group (Fig. 5C). Therefore, the above results provided evidences that *Mettl5*-deficient tumors enhance neoantigen-driven T-cell activation. This expanded TCR landscape may improve recognition capacity for heterogeneous neoantigens generated by aberrant translation, potentially strengthening anti-tumor immunity.

**Fig. 5:**
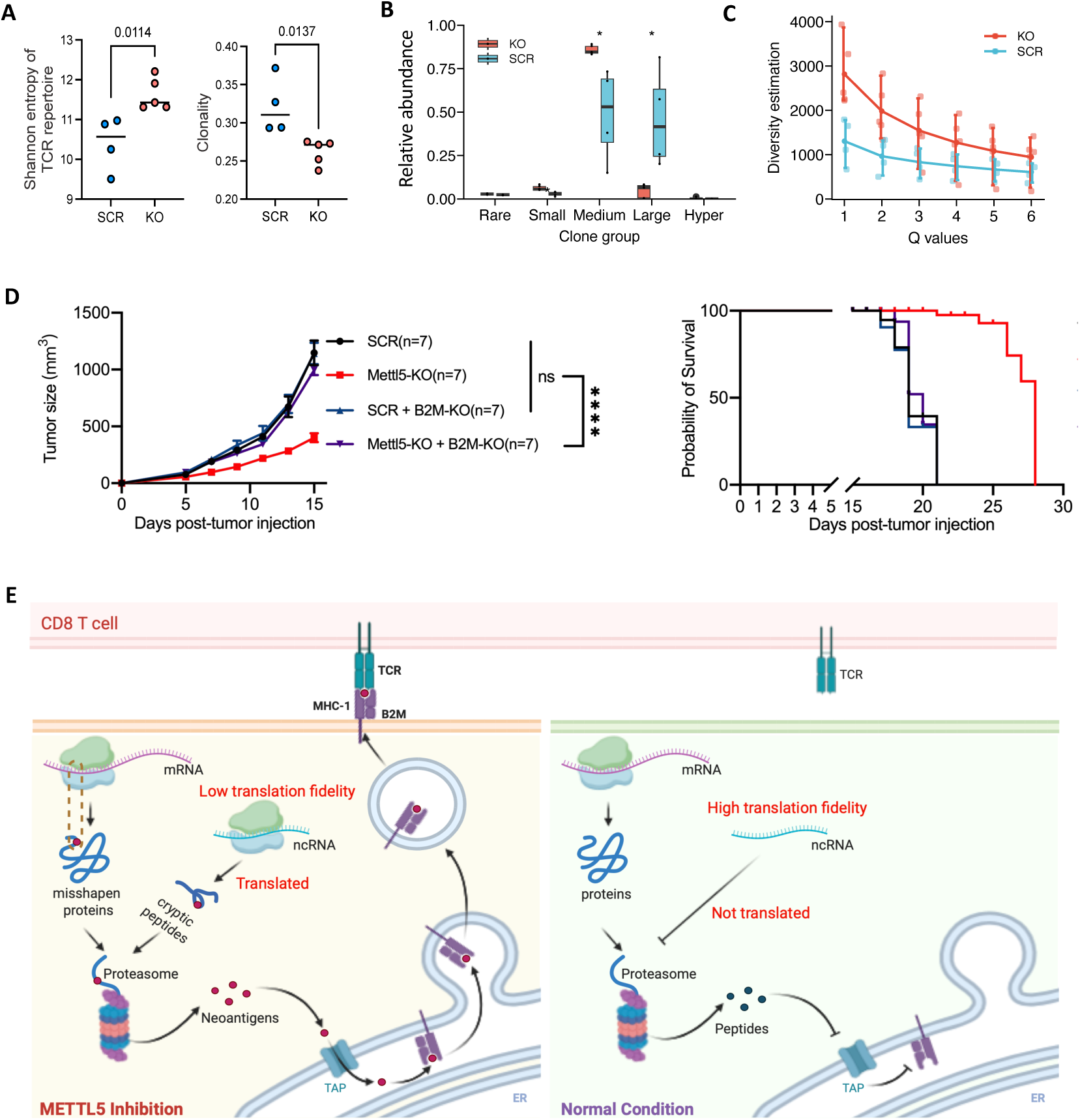
Neoantigens are essential for the anti-tumour immune responses elicited by *Mettl5* depletion. **A**, Scatter plots depicting comparative distributions of TCR repertoire diversity (Shannon entropy) and clonality between *Mettl5*-KO and SCR groups. **B**, Violin plots showing the distribution of clonal expansion levels in *Mettl5*-KO versus SCR groups, with clones categorized by abundance thresholds: Rare (<1e-05), Small (1e-04), Medium (0.001), Large (0.01), and Hyper-expanded (1). **C**, Comparative diversity analysis (Hill numbers q=1–6) between *Mettl5*-KO and SCR groups. **D**, Growth curve of tumors in C57BL/6N mice subcutaneously implanted with SCR, *Mettl5*-KO, SCR plus *B2m*-KO, *Mettl5*-KO plus *B2m*-KO B16-F10 cells. The tumor growth (left) and overall survival (right) were compared. **E**, Working model for the critical role of METTL5 in tumor immunity through neoantigen generation. *Mettl5* deficiency abrogates m⁶A deposition in the ribosomal decoding center, impairing ribosomal function and compromising translation fidelity. These defects lead to aberrant non-canonical translation events that generate immunogenic neoantigens, thereby stimulating anti-tumor immunity. **Note:** Mouse survival curves were plotted using endpoints: death or tumor volume exceeding 2000 mm³. **A, B**, **D**, Student’s t-tests. **A,** Data were represented as mean ± SD. **B**, Data were presented as median (IQR 25-75%). **D** Data were presented as mean ± SEM. \**P*<0.05, \*\*\*\**P*<0.0001, ns: not significant.

Based on our prior evidence that *Mettl5* deficiency enhances MHC-I-restricted neoantigen presentation, we hypothesized that genetic ablation of β2-microglobulin (*B2m*), a critical component of the MHC class I heterodimer, would abolish the antitumor effects observed in METTL5 deficiency tumors. In order to test this hypothesis, *B2m* gene was disrupted using the CRISPR-Cas9 and the knockout efficiency was validated by Western blotting (Fig. S5B, S5C). In syngeneic tumor models, we compared the growth kinetics of tumors derived from *Mettl5-B2m* double-knockout cells and *B2m*-knockout scramble control cells. Both groups exhibited comparable tumor growth rates throughout the experimental period, with no statistically significant differences in tumor volume (Fig. 5D, left). In contrast, the *Mettl5*-KO-only group recapitulated the tumor growth suppression previously observed, confirming the specificity of the antitumor phenotype (Fig. 5D, left). Consistently, the survival analysis also corroborated our hypothesis (Fig. 5D, right). These data definitively established that MHC-I-dependent antigen presentation is indispensable for the antitumor immunity elicited by *Mettl5* depletion.

Taken together, all these findings implicated that the rRNA modification mediated by METTL5 likely plays a critical role in sustaining mRNA translation accuracy. In murine tumor model, loss of *Mettl5* induces mistranslation and promotes the generation of non-canonical translation-derived neoantigens, which in turn boost tumor immunity and represent a novel strategy to enhance immunotherapy (Fig. 5E).

## DISCUSSION

In current study, we identified METTL5 as a critical translation regulator in tumor immune evasion through integrated multi-omics analysis and functional validation. *Mettl5* depletion markedly enhances tumor neoantigen burden by activating translation of noncanonical open reading frame. This mechanism fundamentally differs from traditional mutation-derived neoantigens, as these nuORF-derived neoantigens don’t rely on genomic mutation accumulation but dynamically expand antigenic diversity through disrupting translational fidelity, providing a novel strategy to overcome immunotherapy resistance.

### Molecular mechanism of METTL5 regulating translation fidelity

The dual-luciferase reporter assays and Ribo-Seq analysis indicated that the mRNA translation fidelity is apparently dysregulated upon loss of *Mettl5*. At the molecular level, METTL5-mediated m⁶A1832 modification resides at the h44-h45 helical junction of 18S rRNA, a critical site within the ribosomal decoding center responsible for maintaining codon-anticodon pairing accuracy^53,60,61^. Intriguingly, the m^6^A or m^4^C in the decoding center is dispensable for the ribosome biogensis and cellular viability^53,68^, which implies that they may function primarily in fine-tuning ribosomal activity rather than being essential components. Our findings that METTL5-mediated m⁶A modification ensures translation fidelity in mammals mirror the established role of m⁴C1402 modification in the decoding center of *E. coli*^68,69^, indicating a conserved mechanism for mRNA translation accuracy. However, how these modifications quantitatively affect ribosome function and translational fidelity of mRNA is still not fully understood for now. Future studies should focus on resolving dynamic conformational changes in the ribosomal decoding center following *METTL5* deletion through cryo-electron microscopy to elucidate atomic-level regulatory mechanisms.

### Peptides derived from nuORFs mistranslation as novel sources of neoantigens

In terms of tumor immune microenvironment, *Mettl5*-depleted tumors exhibited a pronounced “hot” phenotype. The flow cytometry analysis and single-cell sequencing revealed increased CD8⁺T cell infiltration density, elevated NK cell proportions, and a T cell receptor (TCR) repertoire with a Shannon diversity index, which further validated the concept that interventing mRNA translation is likely a novel strategy for generating neoantigen and can be utilized for enhancing tumor immuotherapy. Compared to genomic interventions, translational regulation offers unique advantages for neoantigen development: First, the reversibility of translation allows dynamic control of ribosomal function (e.g., modulating methylation levels at the decoding center here) to flexibly adjust translational error rates, avoiding permanent genomic alterations caused by CRISPR or DNA-damaging agents^70–76^. Second, as translational regulation does not directly interfere with DNA sequences, its risks of carcinogenesis and off-target effects are markedly reduced, enhancing clinical safety^70^. Critically, peptide variants generated through translational misreading exhibit high spatiotemporal heterogeneity^73,75^. These transiently existing non-self-epitopes can effectively evade immune editing pressures, breaking immune tolerance induced by fixed mutation-derived epitopes^76^. which sheds light on the future application of neoantigens derived from nuORF mistranslation.

### Targeting ribosome-related factors (RRFs) for enhancing tumor immunity

Recent studies have explored diverse strategies to induce neoantigen generation by targeting tRNA-related pathways, such as inhibiting tRNA modifiers or disrupting tRNA aminoacetylation^76,77^. While these approaches have demonstrated promising preclinical efficacy, their clinical applicability may be limited due to codon-specific restrictions and low diversity of neoantigens produced. In contrast, ribosomal trans-acting factors represent more versatile therapeutic targets. Besides METTL5 reported in current work, multiple ribosome-related factors (RRFs) have been implicated in tumor immunity. Notably, depletion of RPS28 had also been shown to enhance immunogenicity^78^. These findings highlight the broader potential of ribosomal-targeted interventions compared to tRNA-based strategies. On other hand, despite these therapeutic implications, clinical translation of RRFs-targeting strategies faces multiple challenges. The most critical concern is that reduced translational fidelity may lead to aberrant self-antigen production in healthy tissues, potentially triggering autoimmune reactions. Although RRFs are frequently overexpressed in tumors^79,80^ long-term immunological monitoring remains necessary. Nevertheless, these findings highlight RRFs as promising therapeutic targets for cancer immunotherapy. Further investigation is warranted to explore pharmacological and genetic strategies targeting ribosome-related factors to modulate antigen presentation in cancer, particularly tumor immunity contexts.

In summary, our current study not only establishes ribosomal chemical modifications as critical mediators of tumor immune surveillance but also provides experimental evidence for “translational reprogramming” as a novel strategy to overcome current limitations in cancer immunotherapy.

## MATERIALS AND METHODS

### Cell culture

The B16-F10 melanoma cells and 4T1 breast cancer cells were purchased from Wuhan Pricella Biotechnology. All cells were authenticated by STR profiling and testing for mycoplasma contamination was performed with a MycAway one-step mycoplasma detection kit (Yeasen, 40612ES25). All cells were cultured in Dulbecco’s Modified Eagle Medium (DMEM, Gibco) supplemented with 10% fetal bovine serum (FBS, Biological Industries), 1% penicillin-streptomycin (Thermo Fisher Scientific), and 1% SaveIt mycoplasma inhibitor (Hanbio Biotechnology). Cells were maintained in a humidified incubator at 37°C with 5% CO₂ and routinely tested for bacterial and mycoplasma contamination.

### Plasmid construction and Generation of knockout cell

Knockout cells were generated using CRISPR/Cas9 technology. Two independent sgRNAs (targeting *Mettl5* or *B2m* exon 2) were cloned into pL-CRISPR.EFS.GFP plasmid separately. To generate *Mettl5* knockout cells, 2 μg of sgRNA vector that targeting mouse *Mettl5* was transfected to the corresponding cells using Lipofectamine 3000 (Invitrogen). After 2 days’ culture in DMEM medium, cells were screened according to fluorescence and isolated into monoclons by flow cytometry. After 3 weeks’ culture, colonies were screened out and confirmed by sequencing and RT-qPCR. LCMS/MS was also used to further detected the changes of m^6^A modification in SCR and *Mettl5-KO* cells. To generate *Mettl5*-knockout plus *B2m* knockout B16-F10 cells, 2 μg of sgRNA vector that targeting mouse *B2m* was transfected to B16 SCR or *Mettl5-KO* cells using Lipofectamine 3000 (Invitrogen). After 2 days’ culture in DMEM medium, cells were screened according to fluorescence and isolated into monoclons by flow cytometry. After 3 weeks’ culture, colonies were screened out and confirmed by western blot.

sgRNA sequences targeting mouse *Mettl5* locus:

AGCATCGGAGCGGCAATGCTAGG

sgRNA sequences targeting mouse *B2m* locus:

AGTATACTCACGCCACCCACCGG

### Cell proliferation and colony formation

For cell proliferation, B16-F10 (1.5 × 10⁵ cells/well) and 4T1 (0.5 × 10⁵ cells/well) cells were seeded in a 24-well plate on day 0 and counted on day one until day 3 with a cell counter (Thermo Fisher Scientific) to measure cell proliferation. For colony formation, 500 cells were seeded in a 6-well plate and cultured for six days. Cells were then fixed with 4% formaldehyde (Beyotime) and stained with 0.1% crystal violet solution (Solarbio). Next, the number and size of Cell colonies were measured.

### Western blot

For western blotting, a six-well plate of cells at approximately 80-90% confluency was lysed with 100 μL RIPA buffer on ice. Cell lysates were separated by 4-20% SDS-PAGE, and the protein was then transferred to 0.4 μm polyvinylidene fluoride (PVDF, Merck Millipore) membrane. After blocking with 5% milk in TBST, the membranes were then incubated with the corresponding antibodies overnight at 4°C. Membranes were incubated with secondary HRP-conjugated antibody (Proteintech) at room temperature for 2 hours following three times washes with TBST and were washed three times again before detected by the phosphoimager. The corresponding antibodies included Monoclonal rabbit anti-B2m antibody with 1:5000 dilution (Abcam, ab75853) and anti-GAPDH antibody with 1:5000 dilution (Proteintech, 10494-1-AP). Full and uncropped Western blot images are shown in Fig. S6.

### Quantitative Reverse Transcription PCR (RT-qPCR)

Total RNA was isolated from B16-F10 and 4T1 cells using Total RNA Isolation Kit (Vazyme, RC112-01) and reverse-transcribed to cDNA by the use of RT reagent Kit with gDNA Eraser (Takara, RR047A). Primers for quantitative RT-PCR (qPCR) were designed using Primer-BLAST. Reactions were run on the real-time PCR machine (QuantStudio 7 Flex) using TB Green PCR master-mix (Takara, RR430A). The relative gene expression level was calculated with the comparative cycle threshold (CT) method, and the relative number of target transcripts was normalized according to mouse Gapdh gene. Data were plotted using GraphPad Prism 9 (https://www.graphpad.com). Each experiment was performed in triplicate. The oligonucleotide primers used for PCR amplification are listed below:

Mettl5:

Forward primer: 5’-CTGAAGACTGCTTTGGGAATGGC-3’;

Reserve primer: 5’-AATGCTGGTAGATCATATCGAAGC-3’.

Gapdh:

Forward primer: 5’-CATCACTGCCACCCAGAAGACTG-3’;

Reserve primer: 5’-ATGCCAGTGAGCTTCCCGTTCAG-3’.

### Animal Sources

Six-to 8-week-old female C57BL/6N mice were purchased from Beijing Vital River Laboratory Animal Technology Co., Ltd. Six-to 8-week-old female Balb/c mice and 3-to 4-week-old female nude mice were obtained from GemPharmatech. All mice were bred in a specific-pathogen-free (SPF) animal facility with a 12-h light/12-h dark cycle, at 22.5 ± 1°C and 50 ± 5% relative humidity. All animal experiments were approved by the Institutional Animal Care and Use Committee (IACUC) of Southern University of Science and Technology.

### *In vivo* competition assay

For generation of mCherry-labeled SCR and GFP-labeled *Mettl5* knockout KO cells, lentiviral particles were produced by co-transfecting 293T cells with either the GFP or mCherry plasmid along with the packaging plasmids pMD2.G and PsPAX2. B16 SCR cells were transduced with the GFP-expressing lentivirus, while *Mettl5*-KO cells were infected with the mCherry-expressing lentivirus. Following three days of culture, cells were fluorescence-sorted by flow cytometry to isolate stable populations expressing the respective markers. A 1:1 mixture of mCherry-labeled SCR cells and GFP-labeled KO cells was prepared. This mixture was either subcutaneously injected into C57BL/6N mice or co-cultured in vitro. Tumors were harvested 12 days post-injection, and the relative proportions of GFP⁺ (Mettl5-KO) and mCherry⁺ (SCR) cells in both tumors and in vitro cultures were quantified by BD Canto II flow cytometer (Flow Cytometry Shared Facility, SUSTech University).

### Tumour growth in mice

Syngeneic and xenogeneic subcutaneous tumor transplantation assays to evaluate tumorigenicity of mettl5-deficient tumor cells. For syngeneic models, tumor cells were resuspended in PBS (Gibco) and injected subcutaneously into the right dorsal flank of C57BL/6N or Balb/c mice. For xenogeneic models, tumor cells were mixed with an equal volume of Matrigel (Corning) at a 1:1 ratio before subcutaneous injection into nude mice. Tumor volume was measured every other day starting from day 5 or 7 until tumors reached a maximum size of 2000 mm^3^, calculated as length × width^2^. At the experimental endpoint, tumors were harvested and weighed. Mouse survival curves were plotted using endpoints: death or tumor volume exceeding 2000 mm³.

### Preparation of tumor tissue single-cell suspension

At the endpoint of the mouse tumor model, tumors were bluntly dissected intact using surgical scissors and forceps (RWD). After all tumors were stripped, the mass of each tumor sample was recorded and transferred to a 6-cm dish with appropriate HBSS (Gibco). Using curved scissors (RWD), tumors were uniformly minced until passable through a 1-mL pipette tip. Then, collagenase IV (0.5 mg/mL, Sigma) and DNase I (100 U/mL) were added proportionally to the tumor volume. Digestion was performed on a 37℃ shaker at 80 rpm for 45 min. The tumor suspension was filtered through a 70-μm strainer, and single-cell suspension was obtained by centrifugation at 1500 rpm for 10 min at 4℃, followed by three PBS washes. Finally, cells were collected, stained with antibodies, and analyzed by flow cytometry.

### Analysis of tumour-infiltrating lymphocytes

Single-cell suspensions of tumor tissues were prepared and incubated with 50 μL of a solution containing anti-mouse CD16/32 (BioLegend, catalog #101320, diluted 1:50) and anti-mouse Zombie NIR™ Fixable Viability Kit (APC-750, BioLegend, catalog #423105, diluted 1:200) to block non-specific binding and label dead cells. After incubation at 4°C for 10 minutes, a cocktail of surface antibodies for mouse immune cell subset staining was prepared. Cells were stained with the antibody cocktail in 100 μL of tumor staining buffer (BioLegend, #420201) on ice at 4°C for 30 minutes. After incubation, cells were washed twice with staining buffer and resuspended in 900 μL of staining buffer. Counting beads (Invitroge) were added (100 μL per sample) to quantify cell concentrations. Samples were analyzed using a BD Canto II flow cytometer (Flow Cytometry Shared Facility, SUSTech University), with 300,000 CD45^+^ events recorded per sample. Data were analyzed using FlowJo (version 10).

The antibodies used were as follows: anti-mouse CD4 (Pacific Blue, BioLegend, #100531, diluted 1:200); anti-mouse CD45 (Brilliant Violet 510, BioLegend, #103138, diluted 1:100); anti-mouse CD19 (FITC, BioLegend, #115505, diluted 1:200); anti-mouse CD8α (PerCP/Cyanine5.5, BioLegend, #100734, diluted 1:200); anti-mouse NK-1.1 (PE, BioLegend, #108708, diluted 1:200);anti-mouse CD25 (PE/Dazzle 594, BioLegend, #102048, diluted 1:200); anti-mouse TCR β chain (PE/Cyanine7, BioLegend, #109222, diluted 1:200); anti-mouse CD107a (APC, BioLegend, #121615, diluted 1:200); anti-mouse CD11c (APC/Fire™ 750, BioLegend, #117352, diluted 1:200).

### Immunohistochemistry (IHC) detection

IHC was used to evaluate the expression of CD8α in tumor tissues and compare SCR and Mettl5-KO samples. Paraffin-embedded tumor tissues were sectioned into 4-5 μm thick slices and mounted on glass slides. The sections were deparaffinized using xylene and alcohol, followed by antigen retrieval in a citrate buffer. Nonspecific binding was blocked with an appropriate solution, and then a primary antibody specific to the CD8α protein was applied and incubated overnight at 4°C. After washing, a secondary HRP-conjugated antibody was added and incubated for one hour at room temperature. The bound antibodies were visualized using an IHC detection substrate that produced a colorimetric reaction. Counterstaining with hematoxylin was performed to highlight cell nuclei. Images were captured using a microscope, and positive staining cells were quantified within a defined area using image analysis software. Statistical analysis was conducted to assess the differences in the number of stained cells between SCR and KO samples, and the findings were reported with representative images, quantitative data, and a discussion of the results.

### Luciferase Reporter Assay

The dual luciferase reporter vector pmirGLO, which contains both firefly (F-Luc) and Renilla (R-Luc) luciferases, was chosen for meauringing the changes of translational activities. The original pmirGLO plasmids were engineered by introducing a UGA stop codon or frameshift mutations (+1 or −1) into the firefly luciferase gene, respectively. 1 μg of wildtype or mutatant plasmids were transfected into B16-F10 WT or *Mettl5* KO cells. After 48 hours of transfection, luciferase expression was measured using the Dual-Glo Luciferase Assay System (Promega).

### Ribo-seq

Two 15-cm plates of cells with approximately 80-90% confluency was used per group. Cells were incubated with 100 μg/mL Cycloheximide (MCE) for 8 min at 37°C and were then lysed on ice for 10min in 400uL lysis buffer (20 mM Tris–HCl pH 7.5, 150 mM NaCl, 5 mM MgCl2, 1 mM DTT, 100 μg/mL Cycloheximide, 1% Triton X-100). The supernatant was collected via centrifugation at 12 000 g for 10 min at 4 °C, and its’ absorbance at 260 nm was measured. The ribosomes with equal A260 in each group were reserved for RNA sequencing. Added 15U RNase1 (Thermo Fisher Scientific) for each 1OD of lysate, and incubated the reactions at 22°C for 40 min with gentle mixing. Ribosome-protected fragments (RPFs) were collected by MicroSpin S-400 columns (Cytiva) and purified by Zymo Research RNA Clean & Concentrator-25 kit (Zymo). The RPFs were then separated with 15% polyacrylamide TBE–urea gel, excised the gel slices between 17 and 34 nucleotides and recovered RNA from PAGE gel by ZR small-RNA™ PAGE Recovery Kit (Zymo). Purified RNA was treated with PNK(NEB), and the RNA library was prepared by VAHTSTM Small RNA Library Prep Kit for Illumina®(Vazyme). The library concentration was analyzed using the VAHTSTM Library Quantification Kit for Illumina®, and equimolar pools of libraries were sequenced.

### MHC-Associated Peptide Enrichment for Immunopeptidomics Analysis

For MHC-associated peptide isolation, we used two strategies: **1) MHC-I immunoprecipitation (IP):** four 15-cm plates of B16 SCR or Mettl5-KO cells were harvested and subsequently lysed on ice in 8 mL of 1× Lysis buffer (comprising 0.5% NP-40, 50 mM Tris pH 8.0, 150 mM NaCl, and protease inhibitor) for 10 minutes. Following lysis, protein extracts were obtained by centrifugation at 12,000 g for 10 minutes at 4°C. The supernatants were then incubated overnight with anti-mouse MHC-I antibody conjugated to Pierce Protein A/G magnetic beads (Santa Cruz) to facilitate immunoprecipitation. The bead-protein mixture was first treated with 1 mL of Wash Buffer 1 (0.5% NP-40, 50 mM Tris-HCl pH 8.0, 150 mM NaCl, 5 mM EDTA, 100 μM PMSF, and protease inhibitor) to remove nonspecifically bound proteins while simultaneously inhibiting proteolytic degradation. Subsequent washes included 1 mL of Wash Buffer 2 (50 mM Tris-HCl pH 8.0, 150 mM NaCl) to eliminate residual detergent from the first step. High-affinity contaminants were then dissociated using 1 mL of Wash Buffer 3 (50 mM Tris-HCl pH 8.0, 450 mM NaCl), which applied high ionic strength to disrupt non-specific interactions. Finally, a salt-free wash with 1 mL of Wash Buffer 4 (50 mM Tris-HCl pH 8.0) ensured complete removal of residual salts, thereby maintaining compatibility with downstream elution steps. Bound peptides were released by treating the beads with 10% acetic acid (pH ∼3.0), which denatures MHC molecules and facilitates peptide dissociation. The eluate was then filtered through a 3 kDa molecular weight cut-off filter to remove the MHC-I α-chain, B2m protein, and the corresponding MHC antibody. The eluted proteins were ultimately analyzed by HPLC-MS to identify the MHC peptides. **2) Mild acid elution (MAE):** this was performed on six 15 cm dishes of B16 control cells or *Mettl5*-KO cells stimulated with 100IU/mL IFN-γ for 24 h. Cells were treated on ice with pH 3.3 citric acid-phosphate buffer for 5 min to dissociate MHC-antigen peptide complexes and release antigenic peptides. After centrifugation at 10000 g, the supernatant was collected and filtered through a 3 kDa molecular weight cutoff filter. The eluted proteins were analyzed by HPLC-MS for MHC peptides.

### TCR library construction and T-Cell Receptor Repertoire Sequencing

PBMCs were isolated from whole blood of C57BL/6N mice that had been inoculated with either control B16-F10 cells or *Mettl5*-KO cells 12 days prior. DNA was extracted from PBMC using a QIAamp DNA Blood Mini Kit (Qiagen), and the concentration was tested using NanoDrop 2000 spectrophotometer (Thermo Fisher Scientific). The DNA was used as a template for PCR amplification, which was performed to generate the library of TCR. The step 1 PCR amplification protocol was as follows: 95 °C for 5 min; 95 °C for 30 s, 59 °C for 30 s; 72 °C for 1 min for 30 cycles; and 72 °C for 10 min. The step 2 PCR amplification protocol was as follows: 98 °C for 2 min; 98 °C for 30 s, 65 °C for 30 s; 72 °C for 30 s for 10 cycles; and 72 °C for 5 min. The PCR products were purified, and the barcodes were confirmed. The annealed region of these primers was the CDR3 region, as shown by Multiplex PCR Amplifies. The CDR3 regions were sequenced on Illumina NextSeq500 (MyGenostics). According to the sequencing depth, the test covered about 90% of the TCR. The CDR3 regions were identified based on the definition established by the International ImMunoGeneTics (IMGT) (http://www.imgt.org/, accessed on 19 March 2021) collaboration, and the V, D, and J segments contributing to each CDR3 region were identified by a standard algorithm. Data were analysed using ImmunoSEQ Analyzer 3.0, an Adaptive Biotechnologies online analysis platform.

### RNA-seq and Ribo-seq analysis

Adapter sequences were removed from both Ribo-seq and RNA-seq reads using TrimGalore (v0.6.7). For Ribo-seq data, reads shorter than 20 nucleotides or longer than 35 nucleotides were discarded to ensure high-quality ribosome-protected fragment selection. The trimmed reads were then aligned to rRNA and tRNA sequences using Bowtie2 (v2.2.5)^81^. Reads that aligned to rRNA or tRNA were excluded, while the remaining unaligned reads were retained for subsequent analysis. Processed reads were aligned to the reference genome (gencode GRCh38) by the STAR (v2.7.10a)^82^, allowing for at most two mismatches. For Ribo-seq data, quality control and translational profiles was analyzed by RiboseQC. Transcriptome assembly and quantification were performed using StringTie (v2.1.7)^83^. Both read counts of Ribo-seq and RNA-seq were converted to TPM (transcripts per million). Genes with sufficient expression level (Ribo TPM > 1 and RNA TPM > 5) were subjected to further analysis. Translation efficiency (TE) and the differentially-TE genes (DTEGS) were calculated by RiboDiff (v0.2.2)^84^. GO term analysis were performed by Metascape (https://metascape.org/gp/index.html). GraphPad Prismsoftware and R was used for data presentation.

### Hierarchical prediction of translated ORFs across tissues

In our study, we used the RiboCode^85^ algorithm to detect actively translated open reading frames (ORFs) with the parameters set as follows: a minimum fractional threshold of ribosome profiling reads (-f0_percent 0.5), a primary start codon of ATG (--start_codon ATG), alternative start codons of CTG, GTG, and TTG (--alt_start_codons CTG, GTG, TTG), and a minimum amino acid length of 8 (--min-AA-length 8). The identified ORFs were classified into several categories: “annotated” ORFs, which overlap with annotated coding sequences (CDS) and share the same stop codon; “uORFs”, located upstream of the annotated CDS without overlap; “dORFs”, situated downstream of the annotated CDS without overlap; “Overlap_uORFs”, overlapping with the annotated CDS while being upstream; “Overlap_dORFs”, overlapping with the annotated CDS while being downstream; “Internal” ORFs, which are internal to the annotated CDS but in a different reading frame; and “novel” ORFs, originating from non-coding genes or non-coding transcripts of coding genes. Additionally, we generated collapsed BED12 files for annotated ORFs (annotatedORFdb.bed) and non-canonical ORFs (excluding the “annotated” type, nuORFdb.bed), which served as references for subsequent analysis.

### Generating nuORFdb v.1.0

The nuORFdb v.1.0 was constructed following the previous method ^71^. Briefly, to count the number of ribosomal profiling (RPF) reads aligned to ORFs, we first corrected the read positions based on the offset distances between the 5’ end of the reads and the ribosomal A-sites. Subsequently, we employed the “rpfTPM” script from a prior study (available at link) to quantify the number of in-frame reads. This script normalizes the in-frame read counts to transcripts per million (TPM) against the nuORFdb.bed file. ORFs with TPM values 1.5-fold higher in the KO samples compared to SCR samples were designated as “KO-specific nuORFs”. Conversely, ORFs with TPM values 1.5-fold higher in SCR samples relative to KO samples were classified as “SCR-specific nuORFs”.

### Estimation of absolute translation levels

UniProt (reviewed mouse, release 2025_02) and the translated amino acid sequences of larger nuORFs were used as reference databases for protein and peptide identification. Notably, only those nuORFs longer than 8 aa were defined as potential translated regions. The proteomics search was performed using SearchGUI (v4.2.17) (https://github.com/compomics/searchgui) with MS-GF+ and Tide as search engines. Carboxyamidomethylation of cysteine was set as a fixed modification, while oxidation of methionine and proline were set as variable modifications. The enzyme specificity was set to ‘unspecific’ and the precursor mass tolerance was set to 20 ppm Hydrophobicity values for total PSM peptides were predicted with R package protViz (v0.7.9) (https://github.com/cpanse/protViz).

### Statistical analysis

No sample size calculations and blinding were performed. There was no method of randomization. No samples or animals were excluded from analysis. Prism 9 software (GraphPad) was used to analyze data from biological experiments by performing two-tailed unpaired Student’s t-tests. One-way analysis of variance (ANOVA) with Tukey’s test was used to compare multiple groups and two-way ANOVA was used to compare cell proliferation analysis and tumor growth curves. *P* values \**P<*0.05, \*\**P<*0.01, \*\*\**P<*0.001, \*\*\*\**P<*0.0001 were considered statistically significant.

## ACKNOWLEDGEMENTS

We would like to thank all members in Hao Chen’s Lab for their help and advice in experimental design. The authors would also like to acknowledge the technical support from Hua Li and Lin Lin at SUSTech CRF. This work was supported by Center for Computational Science and Engineering at Southern University of Science and Technology.

## AUTHOR CONTRIBUTIONS

H.C. and W.Q.L. designed and conceived the experiments. Y.Y.Z., X.Y.S., R.Q.W., F.L.L. and Y.L.C. performed experiments. Y.C.W. and X.Y.S. analyzed the data. Y.Y.Z., X.Y.S. and H.C. wrote the manuscript. All authors have read and approved the final manuscript.

## DATA AND MATERIALS AVAILABILITY

The raw data of NGS is available in the GEO database under the accession number: GSE299761 (token: ozyvwyqmhnkdfcb). The mass spectrometry proteomics data have been deposited to the ProteomeXchange Consortium via the iProX partner repository with the dataset identifier PXD066586. The proteome datasets can be accessed at https://www.iprox.cn/page/PSV023.html;?url=1753447360479ycXH using password u9OM.

---

**Fig. S1:**
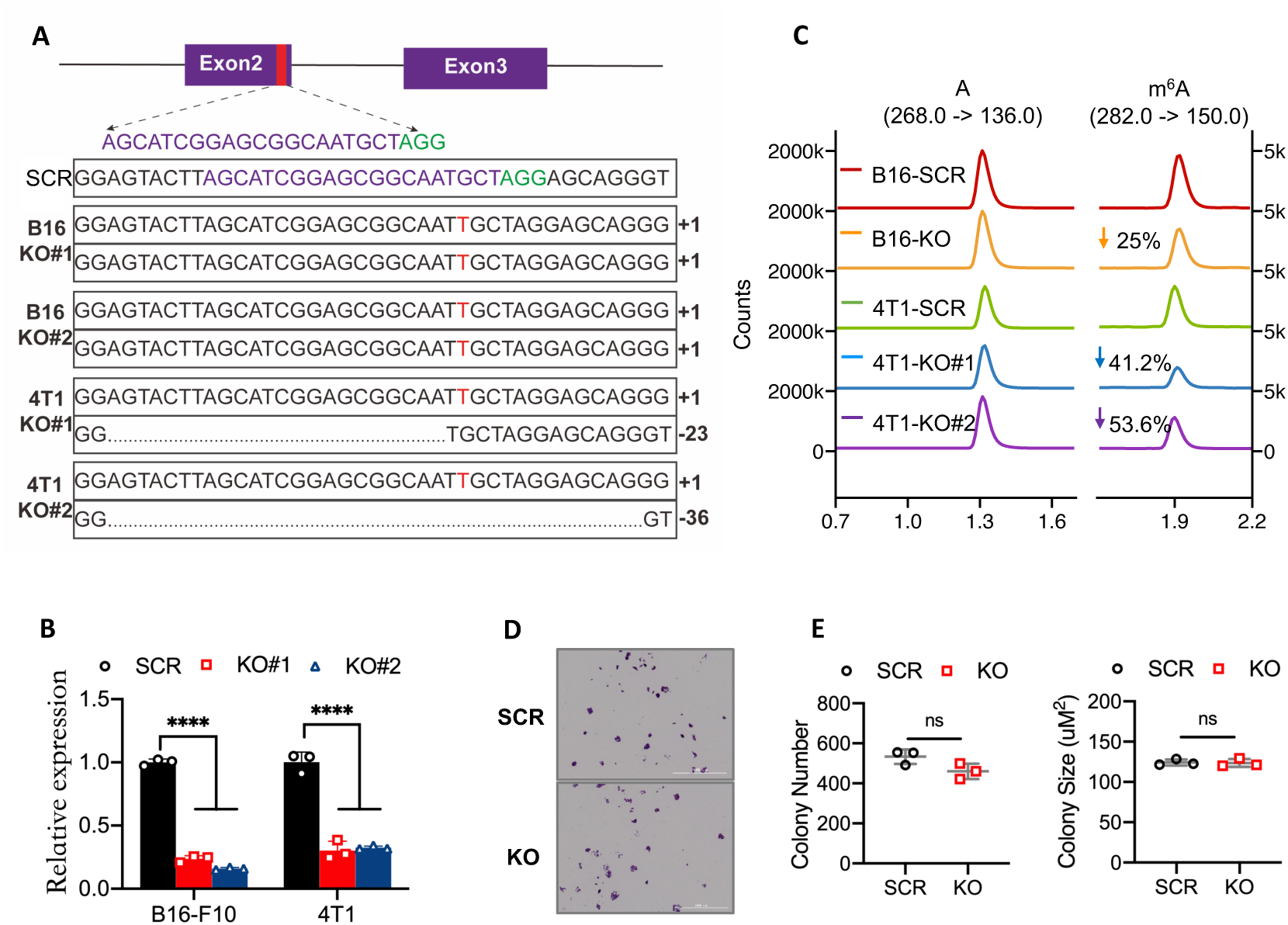
Genetic ablation of *Mettl5* via CRISPR-Cas9 and its effect on tumour-cell growth *in vitro*. **A**, Schematic diagram of sgRNA targeting mouse *Mettl5* locus. Sanger sequencing was used to confirm CRISPR-Cas9-mediated knockout. **B**, Knockout of *Mettl5* in B16-F10 cells and 4T1 cells were verified by Real-time quantitative PCR analysis. **C**, HPLC-MS/MS analysis of m^6^A/A levels in total RNA purified from the SCR and *Mettl5* KO cells. **D**, **E**, Colony-formation assay of WT and *Mettl5* KO B16-F10 cells. 500 cells were seeded in a 6-well plate and cultured for six days, representative pictures (**D**) were taken, and the colony numbers and sizes (**E**) were measured. **Note: B**, One-way ANOVA with Tukey’s test. **E**, Student’s t-tests (right). **B**, **E**, Data were represented as mean ± SD (n=3). \*\*\*\**P*<0.0001, ns: not significant.

**Fig. S2:**
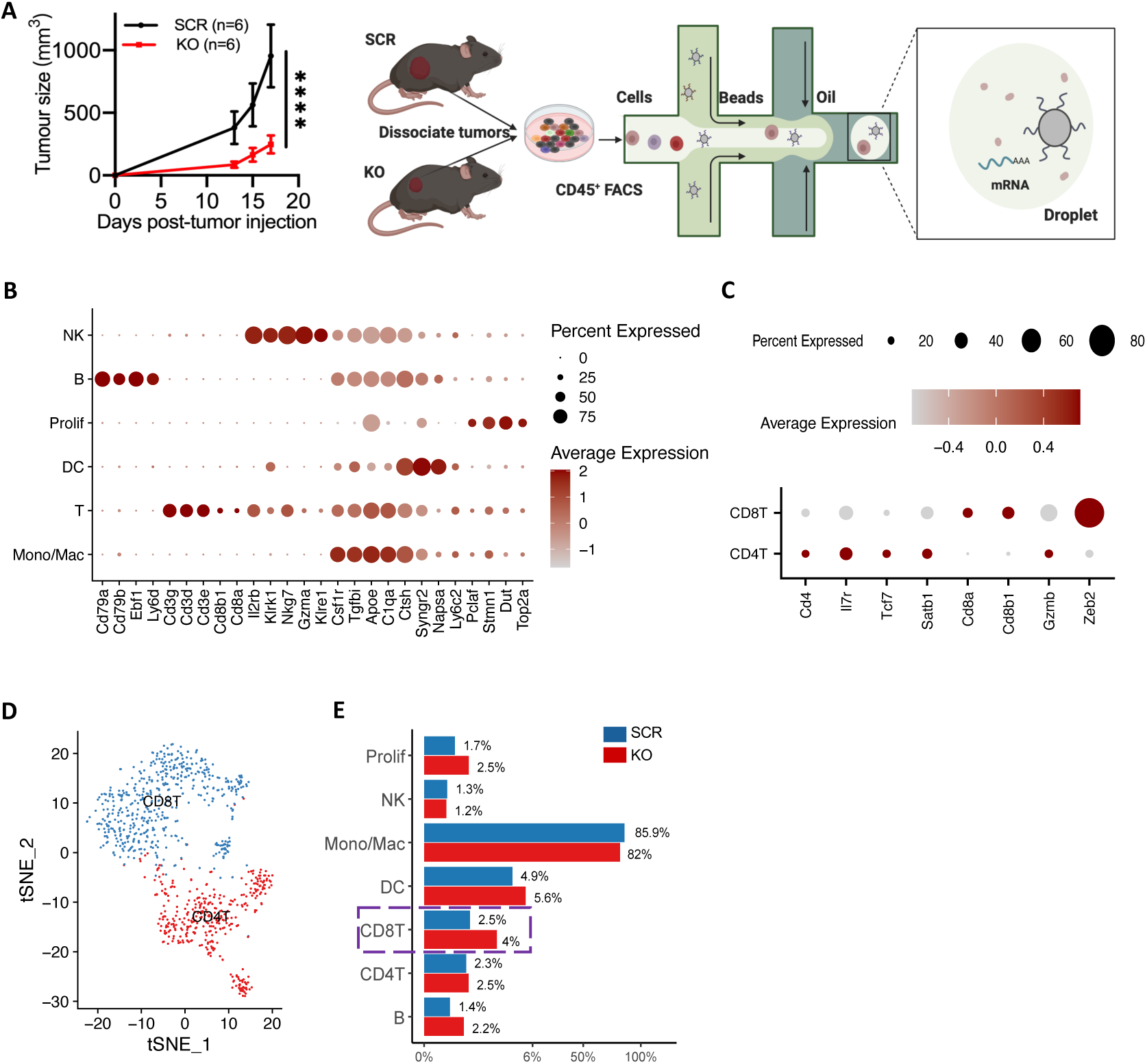
Analysis of the tumor immune cell infiltration by scRNA-seq. **A**, Workflow schematic of single-cell sequencing for tumor tissues. In vivo tumor growth kinetics in C57BL/6 mice inoculated with SCR or *Mettl5*-KO B16-F10 cells. Tumors were excised and dissociated at day 18 post-inoculation, followed by CD45^+^ immune cell enrichment and single-cell sequencing. **B, C,** Dot plot illustrates the clustering analysis results of single-cell sequencing data (**B**). T cells were isolated from the total immune cell population and further stratified into CD8^+^ T cells and CD4^+^ T cell subsets (**C**). Dot color indicates expression level and dot size indicates the proportion of each cell type expressing each gene. **D**, t-SNE plots show T-cell subpopulations derived from tumors, including CD4^+^ T-cell and CD8^+^ T-cell subpopulations. **E**, Bar chart shows the proportion of tumor-infiltrating immune cell populations, with blue representing the *Mettl5*-knockout group and red the control group.

**Fig. S3:**
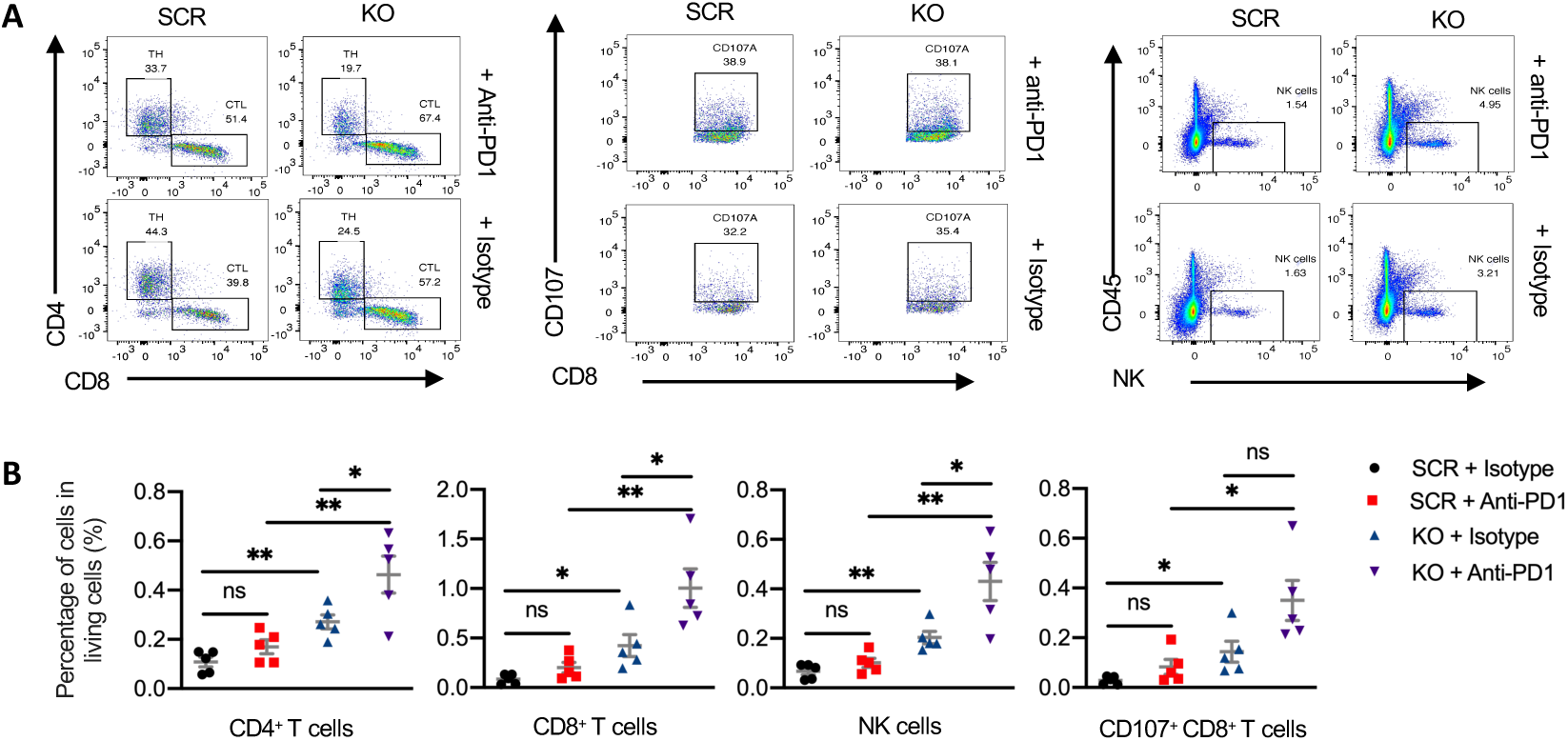
Depletion of *Mettl5* enhances intra-tumoral T-cell infiltration. **A**, **B**, Representative flow cytometry image (**A**) showing the distribution of CD4^+^ T cells, CD8^+^ T cells (left), CD107a^+^ CD8^+^ T cells (middle), and NK cells(right). The relative frequency of immune cells amongst living cells (**B**) were assessed. **Note: B**, Student’s t-tests. Data were represented as mean ± SD (n=5). \**P*<0.05, \*\**P*<0.01, ns: not significant.

**Fig. S4:**
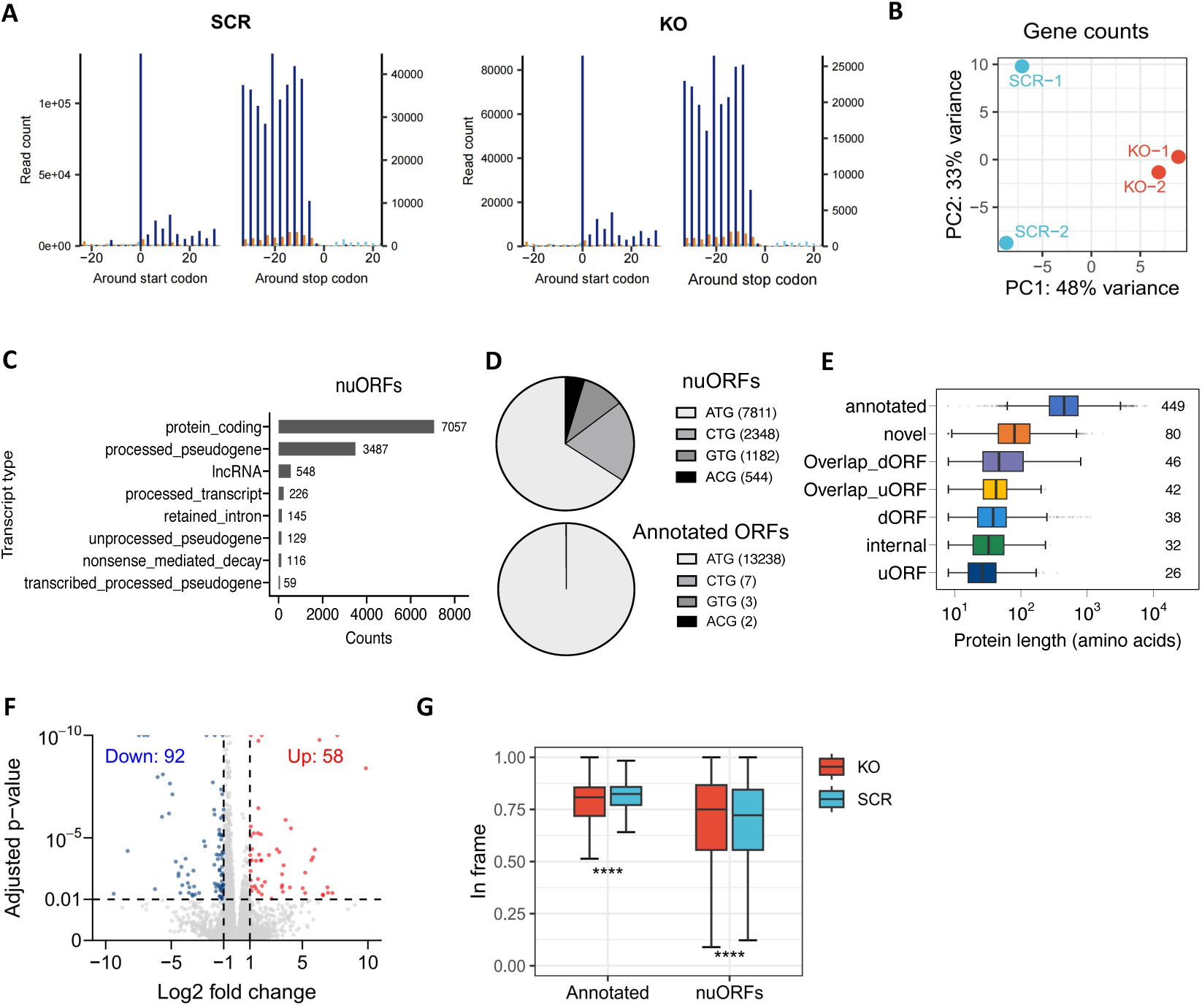
Identification of novel ORFs induced by *Mettl5* deficiency. **A**, Quality control of Ribo-seq data in SCR (left) and *Mettl5* KO (right) cells. **B**, The PCA plot shows the separation between *Mettl5*-KO and SCR groups based on gene counts **C**, Distribution of transcript types for nuORFs identified by Ribo-seq. **D**, Distribution of start codons for nuORFs and annotated ORFs in Mettl5-KO and SCR groups. **E**, Protein length distribution (in amino acids) for different ORF categories in *Mettl5*-KO and SCR groups. **F**, Volcano plot depicting down-regulated (blue) and up-regulated (red) genes identified by Ribo-seq in *Mettl5* KO cells compared with SCR cells. **G**, Distribution of in-frame translation events in *Mettl5*-KO and SCR groups across annotated ORFs and nuORFs. **Note: G**, Student’s t-tests. Data were presented as median (IQR 25-75%). \*\*\*\**P*<0.0001.

**Fig. S5:**
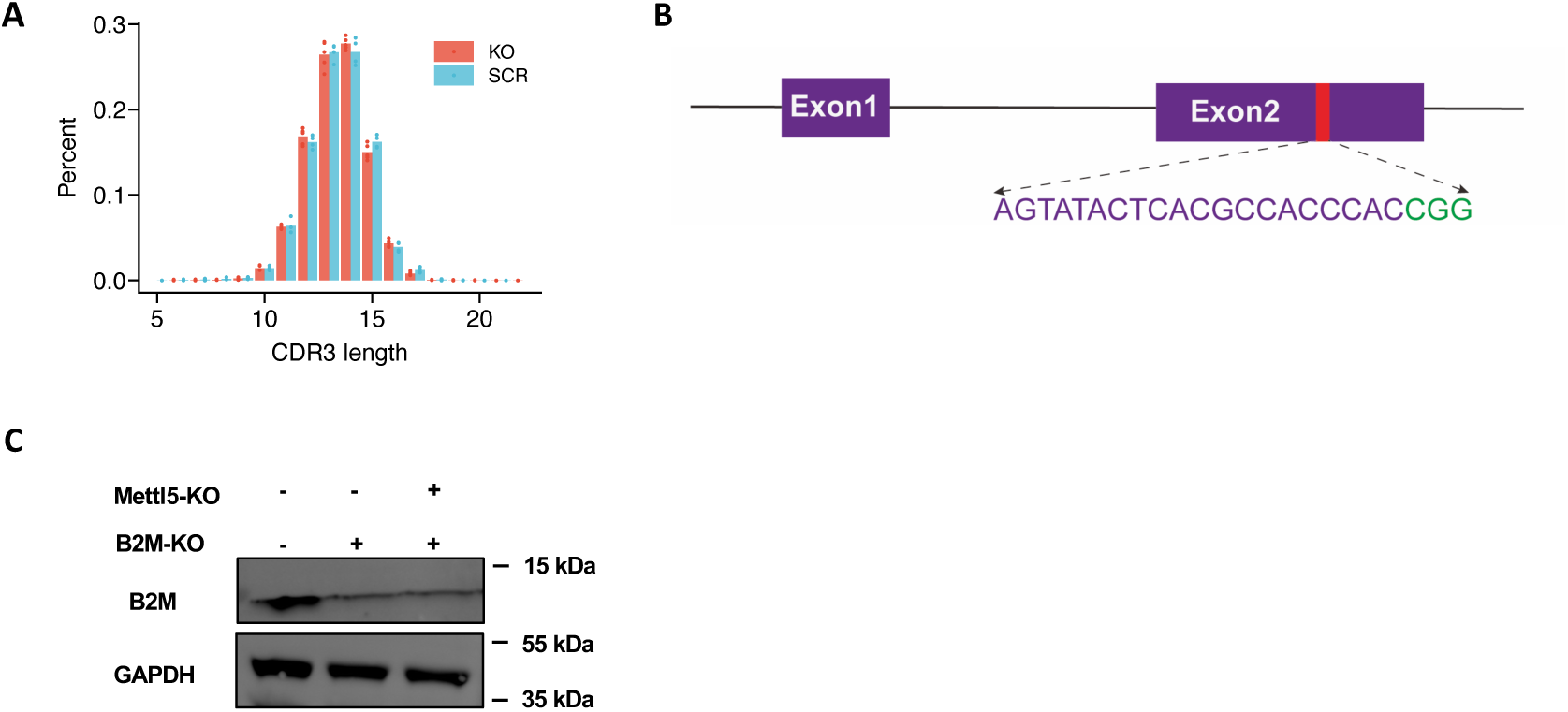
TCR-seq QC and generation of *B2m* knockout cell lines. **A**, Scatter plot from TCR-seq quality control showing no significant differences in CDR3 length distribution between KO and SCR groups. **B**, Schematic diagram of sgRNA targeting mouse *B2m* locus. **C**, Western blotting confirmed the depletion of *B2m* in SCR and *Mettl5*-KO B16-F10 cells. GAPDH was used as internal control.

**Fig. S6:**
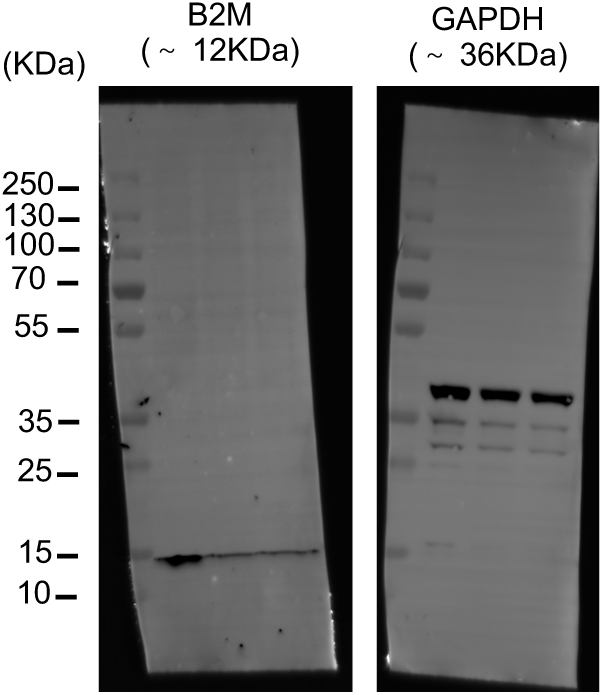
Uncropped original Western blot images.

## REFERENCES

1 Gajewski, T. F. et al. Immune resistance orchestrated by the tumor microenvironment. Immunol Rev 213, 131–145 (2006). 10.1111/j.1600-065X.2006.00442.x

2 Joyce, J. A. & Fearon, D. T. T cell exclusion, immune privilege, and the tumor microenvironment. Science 348, 74–80 (2015). 10.1126/science.aaa6204

3 Hegde, P. S. & Chen, D. S. Top 10 Challenges in Cancer Immunotherapy. Immunity 52, 17–35 (2020). 10.1016/j.immuni.2019.12.011

4 Sharma, P., Hu-Lieskovan, S., Wargo, J. A. & Ribas, A. Primary, Adaptive, and Acquired Resistance to Cancer Immunotherapy. Cell 168, 707–723 (2017). 10.1016/j.cell.2017.01.017

5 Zaretsky, J. M. et al. Mutations Associated with Acquired Resistance to PD-1 Blockade in Melanoma. N Engl J Med 375, 819–829 (2016). 10.1056/NEJMoa1604958

6 McDonald, K. A. et al. Tumor Heterogeneity Correlates with Less Immune Response and Worse Survival in Breast Cancer Patients. Ann Surg Oncol 26, 2191–2199 (2019). 10.1245/s10434-019-07338-3

7 El-Sayes, N., Vito, A. & Mossman, K. Tumor Heterogeneity: A Great Barrier in the Age of Cancer Immunotherapy. Cancers (Basel) 13 (2021). 10.3390/cancers13040806

8 Weiskopf, K. et al. Engineered SIRPalpha variants as immunotherapeutic adjuvants to anticancer antibodies. Science 341, 88–91 (2013). 10.1126/science.1238856

9 Chabner, B. A. & Roberts, T. G., Jr. Timeline: Chemotherapy and the war on cancer. Nat Rev Cancer 5, 65–72 (2005). 10.1038/nrc1529

10 Ahmed, M. M. et al. Harnessing the potential of radiation-induced immune modulation for cancer therapy. Cancer Immunol Res 1, 280–284 (2013). 10.1158/2326-6066.CIR-13-0141

11 Pao, W. et al. Acquired resistance of lung adenocarcinomas to gefitinib or erlotinib is associated with a second mutation in the EGFR kinase domain. PLoS Med 2, e73 (2005). 10.1371/journal.pmed.0020073

12 Oh, T. K. et al. Long-Term Oncologic Outcomes, Opioid Use, and Complications after Esophageal Cancer Surgery. J Clin Med 7 (2018). 10.3390/jcm7020033

13 Nelson, D. B. et al. Surgical margins and risk of local recurrence after wedge resection of colorectal pulmonary metastases. J Thorac Cardiovasc Surg 157, 1648–1655 (2019). 10.1016/j.jtcvs.2018.10.156

14 Sakuishi, K. et al. Targeting Tim-3 and PD-1 pathways to reverse T cell exhaustion and restore anti-tumor immunity. J Exp Med 207, 2187–2194 (2010). 10.1084/jem.20100643

15 Sharma, P. & Allison, J. P. The future of immune checkpoint therapy. Science 348, 56–61 (2015). 10.1126/science.aaa8172

16 Topalian, S. L., Taube, J. M., Anders, R. A. & Pardoll, D. M. Mechanism-driven biomarkers to guide immune checkpoint blockade in cancer therapy. Nat Rev Cancer 16, 275–287 (2016). 10.1038/nrc.2016.36

17 Puig-Saus, C. et al. Neoantigen-targeted CD8(+) T cell responses with PD-1 blockade therapy. Nature 615, 697–704 (2023). 10.1038/s41586-023-05787-1

18 Carreno, B. M. et al. Cancer immunotherapy. A dendritic cell vaccine increases the breadth and diversity of melanoma neoantigen-specific T cells. Science 348, 803–808 (2015). 10.1126/science.aaa3828

19 Borgers, J. S. W. et al. Personalized, autologous neoantigen-specific T cell therapy in metastatic melanoma: a phase 1 trial. Nat Med 31, 881–893 (2025). 10.1038/s41591-024-03418-4

20 Braun, D. A. et al. A neoantigen vaccine generates antitumour immunity in renal cell carcinoma. Nature 639, 474–482 (2025). 10.1038/s41586-024-08507-5

21 Danelli, L. Personalized neoantigen therapy for melanoma immunotherapy. Nat Cancer 5, 1783 (2024). 10.1038/s43018-024-00850-w

22 Ott, P. A. et al. An immunogenic personal neoantigen vaccine for patients with melanoma. Nature 547, 217–221 (2017). 10.1038/nature22991

23 Ott, P. A. et al. A Phase Ib Trial of Personalized Neoantigen Therapy Plus Anti-PD-1 in Patients with Advanced Melanoma, Non-small Cell Lung Cancer, or Bladder Cancer. Cell 183, 347–362 e324 (2020). 10.1016/j.cell.2020.08.053

24 Parkhurst, M. et al. Adoptive transfer of personalized neoantigen-reactive TCR-transduced T cells in metastatic colorectal cancer: phase 2 trial interim results. Nat Med 30, 2586–2595 (2024). 10.1038/s41591-024-03109-0

25 Wang, M. et al. Humanized dual-targeting antibody-drug conjugates specific to MET and RON receptors as a pharmaceutical strategy for the treatment of cancers exhibiting phenotypic heterogeneity. Acta Pharmacol Sin (2025). 10.1038/s41401-024-01458-7

26 Lim, Z. F. & Ma, P. C. Emerging insights of tumor heterogeneity and drug resistance mechanisms in lung cancer targeted therapy. J Hematol Oncol 12, 134 (2019). 10.1186/s13045-019-0818-2

27 Dash, P. et al. Quantifiable predictive features define epitope-specific T cell receptor repertoires. Nature 547, 89–93 (2017). 10.1038/nature22383

28 Tran, E. et al. Immunogenicity of somatic mutations in human gastrointestinal cancers. Science 350, 1387–1390 (2015). 10.1126/science.aad1253

29 Tran, E. et al. Cancer immunotherapy based on mutation-specific CD4+ T cells in a patient with epithelial cancer. Science 344, 641–645 (2014). 10.1126/science.1251102

30 Lu, Y. C. et al. Mutated PPP1R3B is recognized by T cells used to treat a melanoma patient who experienced a durable complete tumor regression. J Immunol 190, 6034–6042 (2013). 10.4049/jimmunol.1202830

31 Alban, T. J. et al. Neoantigen immunogenicity landscapes and evolution of tumor ecosystems during immunotherapy with nivolumab. Nat Med 30, 3209–3222 (2024). 10.1038/s41591-024-03240-y

32 Lopez, J. et al. Autogene cevumeran with or without atezolizumab in advanced solid tumors: a phase 1 trial. Nat Med 31, 152–164 (2025). 10.1038/s41591-024-03334-7

33 Huber, F. et al. A comprehensive proteogenomic pipeline for neoantigen discovery to advance personalized cancer immunotherapy. Nat Biotechnol (2024). 10.1038/s41587-024-02420-y

34 Liu, C. et al. KRAS-G12D mutation drives immune suppression and the primary resistance of anti-PD-1/PD-L1 immunotherapy in non-small cell lung cancer. Cancer Commun (Lond) 42, 828–847 (2022). 10.1002/cac2.12327

35 Platten, M. et al. A vaccine targeting mutant IDH1 in newly diagnosed glioma. Nature 592, 463–468 (2021). 10.1038/s41586-021-03363-z

36 Schumacher, T. et al. A vaccine targeting mutant IDH1 induces antitumour immunity. Nature 512, 324–327 (2014). 10.1038/nature13387

37 Hsiue, E. H. et al. Targeting a neoantigen derived from a common TP53 mutation. Science 371 (2021). 10.1126/science.abc8697

38 Rojas, L. A. et al. Personalized RNA neoantigen vaccines stimulate T cells in pancreatic cancer. Nature 618, 144–150 (2023). 10.1038/s41586-023-06063-y

39 Keskin, D. B. et al. Neoantigen vaccine generates intratumoral T cell responses in phase Ib glioblastoma trial. Nature 565, 234–239 (2019). 10.1038/s41586-018-0792-9

40 Hu, Z. et al. Personal neoantigen vaccines induce persistent memory T cell responses and epitope spreading in patients with melanoma. Nat Med 27, 515–525 (2021). 10.1038/s41591-020-01206-4

41 Apoorvi, T., Yury, P., Iryna, V. & Michelle, K. Phosphopeptide Neoantigens as Emerging Targets in Cancer Immunotherapy. J Cancer Immunol (Wilmington) 6, 135–147 (2024). 10.33696/cancerimmunol.6.094

42 Gubin, M. M. et al. Checkpoint blockade cancer immunotherapy targets tumour-specific mutant antigens. Nature 515, 577–581 (2014). 10.1038/nature13988

43 McGranahan, N. et al. Clonal neoantigens elicit T cell immunoreactivity and sensitivity to immune checkpoint blockade. Science 351, 1463–1469 (2016). 10.1126/science.aaf1490

44 Schumacher, T. N. & Schreiber, R. D. Neoantigens in cancer immunotherapy. Science 348, 69–74 (2015). 10.1126/science.aaa4971

45 Yadav, M. et al. Predicting immunogenic tumour mutations by combining mass spectrometry and exome sequencing. Nature 515, 572–576 (2014). 10.1038/nature14001

46 Rosenberg, S. A. & Restifo, N. P. Adoptive cell transfer as personalized immunotherapy for human cancer. Science 348, 62–68 (2015). 10.1126/science.aaa4967

47 Snyder, A. et al. Genetic basis for clinical response to CTLA-4 blockade in melanoma. N Engl J Med 371, 2189–2199 (2014). 10.1056/NEJMoa1406498

48 Jackson, S. P. & Bartek, J. The DNA-damage response in human biology and disease. Nature 461, 1071–1078 (2009). 10.1038/nature08467

49 Brown, J. S., O’Carrigan, B., Jackson, S. P. & Yap, T. A. Targeting DNA Repair in Cancer: Beyond PARP Inhibitors. Cancer Discov 7, 20–37 (2017). 10.1158/2159-8290.CD-16-0860

50 Olive, J. F. et al. Accounting for tumor heterogeneity when using CRISPR-Cas9 for cancer progression and drug sensitivity studies. PLoS One 13, e0198790 (2018). 10.1371/journal.pone.0198790

51 Parsels, L. A. et al. Translation of DNA Damage Response Inhibitors as Chemoradiation Sensitizers From the Laboratory to the Clinic. Int J Radiat Oncol Biol Phys 111, e38–e53 (2021). 10.1016/j.ijrobp.2021.07.1708

52 Kovacs, S. A., Fekete, J. T. & Gyorffy, B. Predictive biomarkers of immunotherapy response with pharmacological applications in solid tumors. Acta Pharmacol Sin 44, 1879–1889 (2023). 10.1038/s41401-023-01079-6

53 van Tran, N. et al. The human 18S rRNA m6A methyltransferase METTL5 is stabilized by TRMT112. Nucleic Acids Res 47, 7719–7733 (2019). 10.1093/nar/gkz619

54 Chen, H. et al. METTL5, an 18S rRNA-specific m^6^A methyltransferase, modulates expression of stress response genes. bioRxiv, 2020.2004.2027.064162 (2020). 10.1101/2020.04.27.064162

55 Sepich-Poore, C. et al. The METTL5-TRMT112 N^6^-methyladenosine methyltransferase complex regulates mRNA translation via 18S rRNA methylation. Journal of Biological Chemistry 298 (2022). 10.1016/j.jbc.2022.101590

56 Golstein, P. & Griffiths, G. M. An early history of T cell-mediated cytotoxicity. Nat Rev Immunol 18, 527–535 (2018). 10.1038/s41577-018-0009-3

57 Aktas, E., Kucuksezer, U. C., Bilgic, S., Erten, G. & Deniz, G. Relationship between CD107a expression and cytotoxic activity. Cell Immunol 254, 149–154 (2009). 10.1016/j.cellimm.2008.08.007

58 Delaunay, S., Helm, M. & Frye, M. RNA modifications in physiology and disease: towards clinical applications. Nat Rev Genet 25, 104–122 (2024). 10.1038/s41576-023-00645-2

59 Milenkovic, I. & Novoa, E. M. Dynamic rRNA modifications as a source of ribosome heterogeneity. Trends Cell Biol 35, 604–614 (2025). 10.1016/j.tcb.2024.10.001

60 Xing, M. et al. The 18S rRNA m(6) A methyltransferase METTL5 promotes mouse embryonic stem cell differentiation. EMBO Rep 21, e49863 (2020). 10.15252/embr.201949863

61 Rong, B. et al. Ribosome 18S m(6)A Methyltransferase METTL5 Promotes Translation Initiation and Breast Cancer Cell Growth. Cell Rep 33, 108544 (2020). 10.1016/j.celrep.2020.108544

62 Li, X., Yang, G., Ma, L., Tang, B. & Tao, T. N(6)-methyladenosine (m(6)A) writer METTL5 represses the ferroptosis and antitumor immunity of gastric cancer. Cell Death Discov 10, 402 (2024). 10.1038/s41420-024-02166-1

63 Khatter, H., Myasnikov, A. G., Natchiar, S. K. & Klaholz, B. P. Structure of the human 80S ribosome. Nature 520, 640–645 (2015). 10.1038/nature14427

64 Zhu, X., Cruz, V. E., Zhang, H., Erzberger, J. P. & Mendell, J. T. Specific tRNAs promote mRNA decay by recruiting the CCR4-NOT complex to translating ribosomes. Science 386, eadq8587 (2024). 10.1126/science.adq8587

65 Milicevic, N., Jenner, L., Myasnikov, A., Yusupov, M. & Yusupova, G. mRNA reading frame maintenance during eukaryotic ribosome translocation. Nature 625, 393–400 (2024). 10.1038/s41586-023-06780-4

66 Djumagulov, M. et al. Accuracy mechanism of eukaryotic ribosome translocation. Nature 600, 543–546 (2021). 10.1038/s41586-021-04131-9

67 Holm, M. et al. mRNA decoding in human is kinetically and structurally distinct from bacteria. Nature 617, 200–207 (2023). 10.1038/s41586-023-05908-w

68 Kimura, S. & Suzuki, T. Fine-tuning of the ribosomal decoding center by conserved methyl-modifications in the Escherichia coli 16S rRNA. Nucleic Acids Res 38, 1341–1352 (2010). 10.1093/nar/gkp1073

69 Baudin-Baillieu, A. et al. Nucleotide modifications in three functionally important regions of the Saccharomyces cerevisiae ribosome affect translation accuracy. Nucleic Acids Res 37, 7665–7677 (2009). 10.1093/nar/gkp816

70 Laumont, C. M. et al. Noncoding regions are the main source of targetable tumor-specific antigens. Sci Transl Med 10 (2018). 10.1126/scitranslmed.aau5516

71 Ouspenskaia, T. et al. Unannotated proteins expand the MHC-I-restricted immunopeptidome in cancer. Nat Biotechnol 40, 209–217 (2022). 10.1038/s41587-021-01021-3

72 Laumont, C. M. et al. Global proteogenomic analysis of human MHC class I-associated peptides derived from non-canonical reading frames. Nat Commun 7, 10238 (2016). 10.1038/ncomms10238

73 Mohler, K. & Ibba, M. Translational fidelity and mistranslation in the cellular response to stress. Nat Microbiol 2, 17117 (2017). 10.1038/nmicrobiol.2017.117

74 Chong, C. et al. Integrated proteogenomic deep sequencing and analytics accurately identify non-canonical peptides in tumor immunopeptidomes. Nat Commun 11, 1293 (2020). 10.1038/s41467-020-14968-9

75 Ruiz Cuevas, M. V., et al. Most non-canonical proteins uniquely populate the proteome or immunopeptidome. Cell Rep 34, 108815 (2021). 10.1016/j.celrep.2021.108815

76 Weller, C. et al. Translation dysregulation in cancer as a source for targetable antigens. Cancer Cell 43, 823–840 e818 (2025). 10.1016/j.ccell.2025.03.003

77 Pataskar, A. et al. Tryptophan depletion results in tryptophan-to-phenylalanine substitutants. Nature 603, 721–727 (2022). 10.1038/s41586-022-04499-2

78 Wei, J. et al. Ribosomal Proteins Regulate MHC Class I Peptide Generation for Immunosurveillance. Mol Cell 73, 1162–1173 e1165 (2019). 10.1016/j.molcel.2018.12.020

79 Ramalho, S., Dopler, A. & Faller, W. J. Ribosome specialization in cancer: a spotlight on ribosomal proteins. NAR Cancer 6, zcae029 (2024). 10.1093/narcan/zcae029

80 Truitt, M. L. & Ruggero, D. New frontiers in translational control of the cancer genome. Nat Rev Cancer 16, 288–304 (2016). 10.1038/nrc.2016.27

81 Langmead, B., Trapnell, C., Pop, M. & Salzberg, S. L. Ultrafast and memory-efficient alignment of short DNA sequences to the human genome. Genome Biol 10, R25 (2009). 10.1186/gb-2009-10-3-r25

82 Dobin, A. et al. STAR: ultrafast universal RNA-seq aligner. Bioinformatics 29, 15–21 (2013). 10.1093/bioinformatics/bts635

83 Pertea, M. et al. StringTie enables improved reconstruction of a transcriptome from RNA-seq reads. Nat Biotechnol 33, 290–295 (2015). 10.1038/nbt.3122

84 Zhong, Y. et al. RiboDiff: detecting changes of mRNA translation efficiency from ribosome footprints. Bioinformatics 33, 139–141 (2017). 10.1093/bioinformatics/btw585

85 Xiao, Z. et al. De novo annotation and characterization of the translatome with ribosome profiling data. Nucleic Acids Res 46, e61 (2018). 10.1093/nar/gky179

